# Electrocorticographic and Astrocytic Signatures of Stearoyl-CoA Desaturase Inhibition in the Triple Transgenic Mouse Model of Alzheimer’s Disease

**DOI:** 10.1101/2024.03.01.582986

**Authors:** Audrey Hector, Maria João da Costa Caiado, Tanya Leduc, Benoît Delignat-Lavaud, Julien Dufort-Gervais, Clément Bourguignon, Jean-Marc Lina, Karl Fernandes, Jonathan Brouillette, Valérie Mongrain

## Abstract

The symptomatology of Alzheimer’s disease (AD) includes cognitive deficits and sleep disturbances. Recent findings suggest the involvement of dysfunctions in lipid metabolism, such as oleic acid build-up, in the brain of AD patients and animal models. In addition, the inhibition of stearoyl-CoA desaturase (SCD), a lipid-converting enzyme, was shown to restore memory in triple transgenic (3xTg)-AD mice. In the brain, astrocytes regulate the synthesis of specific lipids. Alterations in astrocytes and their function were reported in AD patients and animal models, and astrocytes have been implicated in the regulation of sleep. However, the relationship between sleep disturbances, astrocytes and lipid metabolism remains to be explored in AD. This project thus aimed at assessing whether the inhibition of SCD restores sleep in 3xTg-AD mice, and whether this associated with modifications in astrocytic function. Wild-type (WT) and 3xTg-AD female mice (4-months old) received intracerebroventricular infusion of a SCD inhibitor (SCDi) or vehicle for 28 days, and a 24-hour electrocorticographic (ECoG) recording was conducted post-treatment. Post-mortem brain slices were stained for the astrocytic markers glial fibrillary acidic protein (GFAP) and 10-formyltetrahydrofolate dehydrogenase (ALDH1L1) to perform cell counting and/or morphological evaluation in the hippocampus, lateral hypothalamus and thalamus. The results indicate that the reduced time spent awake and increased time spent in slow wave sleep (SWS) in 3xTg-AD mice was not restored by the SCDi treatment. Similar observations were made concerning the increased number of wake and SWS bouts in 3xTg-AD mice. Rhythmic and scale-free ECoG activity were markedly altered in 3xTg-AD mice for all wake/sleep states, and SCDi significantly altered these phenotypes in a different manner in mutant mice in comparison to WT mice. GFAP- and ALDH1L1-positive cell densities were elevated in the hippocampus and lateral hypothalamus/thalamus of 3x-Tg-AD mice, respectively, and SCDi rescued the increase in the CA1 region in particular. Overall, these findings suggest that the multiple wake/sleep alterations in 3xTg-AD mice are not substantially restored by targeting lipid metabolism using SCD inhibition, at least for the targeted age window, but that this treatment can revert hippocampal changes in astrocytes. This work will benefit the understanding of the pathophysiology related to AD and associated sleep disturbances.

## INTRODUCTION

Alzheimer’s disease (AD), the most common type of dementia, is characterized by a cognitive decline affecting memory and by sleep disturbances (Wang and Holtzman, 2020; Zhang et al., 2022). In fact, sleep disturbances, such as insomnia and fragmented sleep, are proposed to be a major risk factor for the development of cognitive deficits and AD (Shi et al., 2018; Wang and Holtzman, 2020). Multiple alterations in sleep architecture and/or electrocorticographic (ECoG) activity during sleep are also reported in animal models of AD (Dufort-Gervais et al., 2019; Kent et al., 2019; Kosel et al., 2020; Wang and Holtzman, 2020; Hector et al., 2023), including in the widely used triple transgenic (3xTg)-AD mouse model (Benthem et al., 2020; Saber et al., 2021). For instance, 3xTg-AD mice show a reduced number of very long sleep bouts and more sleep during the dark/active period (Saber et al., 2021), together with modifications in the 24-hour rhythm of wheel-running activity (Sterniczuk et al., 2010; Adler et al., 2019).

Patients with AD, together with animal models, were also reported to have alterations in lipids and their metabolism (Chan et al., 2012; Whiley et al., 2014; Hamilton et al., 2015; Chatterjee et al., 2016). Indeed, specific phospholipids were found to be increased in different brain regions and decreased in the plasma of AD patients (Chan et al., 2012; Whiley et al., 2014), and an elevated level of monounsaturated fatty acids (MUFAs) was reported in the AD brain (Fraser et al., 2010; Astarita et al., 2011). In addition, patients suffering from AD were shown to have a higher mRNA expression of stearoyl-CoA desaturase (SCD), the rate-limiting enzyme in the biosynthesis of MUFAs (Astarita et al., 2011). Interestingly, inhibiting SCD was shown to improve different AD-related phenotypes in the 3xTg-AD mouse model, such as the elevated MUFA level and the deficit in spatial memory (Hamilton et al., 2015, 2022). The potential for this intervention to also improve wake/sleep alterations remains to be unveiled.

Neuroinflammation and alterations in glial functions are expected to contribute to the pathophysiology of AD (Heneka et al., 2015). In 3xTg-AD mice, modifications in the well-recognized astrocytic marker glial fibrillary acidic protein (GFAP) have been reported for several brain regions (Yeh et al., 2011; González-Molina et al., 2021). Importantly, astrocytes play a crucial role in lipid synthesis, distribution, and storage (Lee et al., 2021; Smolič et al., 2021), and dysregulation of lipid metabolism in astrocytes can contribute to brain diseases and neurodegeneration (Chen et al., 2023; Mi et al., 2023). Astrocytes have also been implicated in the regulation of ECoG activity during different sleep states (Halassa et al. 2009; Foley et al., 2017), and a higher expression of activated astrocyte marker genes in the prefrontal cortex was reported to be associated with indications of sleep disturbances and cognitive decline in humans (Wu et al., 2023). Moreover, optogenetic activation of astrocytes was recently shown to rescue sleep oscillations and fear memory in a mouse model of AD (Lee et al., 2023). Therefore, a lipid metabolism-targeting pharmacological intervention for AD could impact sleep via an effect on astrocytic function in key brain regions.

We here aimed to test the hypothesis that the inhibition of SCD can rescue wake/sleep alterations in 3xTg-AD mice via a mechanism involving astrocytes. To test this hypothesis, we used an extensive wake/sleep phenotyping comprising advanced ECoG signal analyses to interrogate variables related to the architecture of vigilance states (i.e., time spent in wakefulness, slow wave sleep [SWS], and paradoxical sleep [PS]; alternations between these states), and to the quality of the ECoG signal during these states. More precisely, standard spectral analyses of the ECoG (e.g., dynamics of slow-wave activity [SWA] during SWS) were combined to an analysis of scale-free activity using Wavelet-Leaders to quantify rhythmic and arrhythmic ECoG properties, which are both highly relevant to AD (Averna et al., 2023; Wang and Holtzman, 2020). This was done together with an evaluation of astrocytes using measurements of GFAP and aldehyde dehydrogenase 1 family member L1 (ALDH1L1; also known as 10-formyltetrahydrofolate dehydrogenase) in hippocampal regions, in the lateral hypothalamus (LH), and in thalamic nuclei. A 28-day treatment was used to align with the previous literature showing positive effects of this duration of SCD inhibition on neurogenesis and memory in 3xTg-AD mice (Hamilton et al., 2015, 2022). Our findings reveal that the specific ECoG signatures of SCD inhibition and of the 3xTg mouse model are mostly independent, but that inhibiting SCD shows the potential to revert astrocytic alterations in the hippocampus.

## MATERIAL AND METHODS

### Animals and protocol

Four-month old female 3xTg-AD mice (B6;129-Tg(APPSwe,tauP301L)1Lfa *Psen1^tm1Mpm^*/Mmjax, Jackson Laboratory MMRRC strain #034830-JAX) and WT (B6129SF2/J, Jackson Laboratory strain #101045) were used in this study. The 3xTg-AD mouse model features three mutations associated with familial AD, namely APP Swedish, MAPT P301L, and PSEN1 M146V (Oddo et al., 2003). It presents an age-dependent progressive AD-like neurodegeneration, including amyloid β deposition in different brain areas and synaptic dysfunctions (Belfiore et al., 2019; Oddo et al., 2003). Importantly, 3xTg-AD females show worse phenotypes than males regarding Aβ pathology (Hirata-Fukae et al., 2008), which is in line with the higher risk of developing AD in women (Lautenschlager et al., 1996). Mice were maintained at the CRCHUM, and sent to the Hôpital du Sacré-Cœur de Montréal (CIUSSS-NIM) for experiments where they were individually housed under a 12-hour light/12-hour dark cycle at 23-25°C room temperature, with free access to food and water, for the full duration of the experiment.

Mice of each genotype were assigned to one of two treatment groups, namely treatment with SCDi or with vehicle, for a total of four experimental groups: WT mice with vehicle (n = 12; age and weight at the time of surgery ± standard error of the mean: 19.3 ± 0.3 weeks, 20.5 ± 0.5 g), WT with SCDi (n = 12; 18.8 ± 0.3 weeks, 21.5 ± 0.7 g), 3xTg mice with vehicle (n = 11; 19.1 ± 0.3 weeks, 25.2 ± 1.1 g), and 3xTg with SCDi (n = 10; 19.0 ± 0.2 weeks, 25.8 ± 1.2 g). Mice were implanted with osmotic minipump for treatment infusion and with electrodes for electrocorticography (ECoG) and electromyography (EMG) during the same surgical procedure (described below; Fig. 1A). Following 28 days of treatment, ECoG/EMG signals were recorded for 24 hours starting at light onset (= Zeitgeber time 0: ZT0). Three days after, animals were sacrificed by cervical dislocation (Fig. 1A), and brains were immediately sampled, separated into left and right hemispheres along the middle sagittal line, and frozen on dry ice. All procedures with animals were conducted according to the guidelines of the Canadian Council on Animal Care, and the protocol was approved by the *Comité d’éthique de l’expérimentation animale* of the CIUSSS-NIM. In addition, all analyses were carried out by experimenters who were blind to animals’ group.

**Fig. 1.**
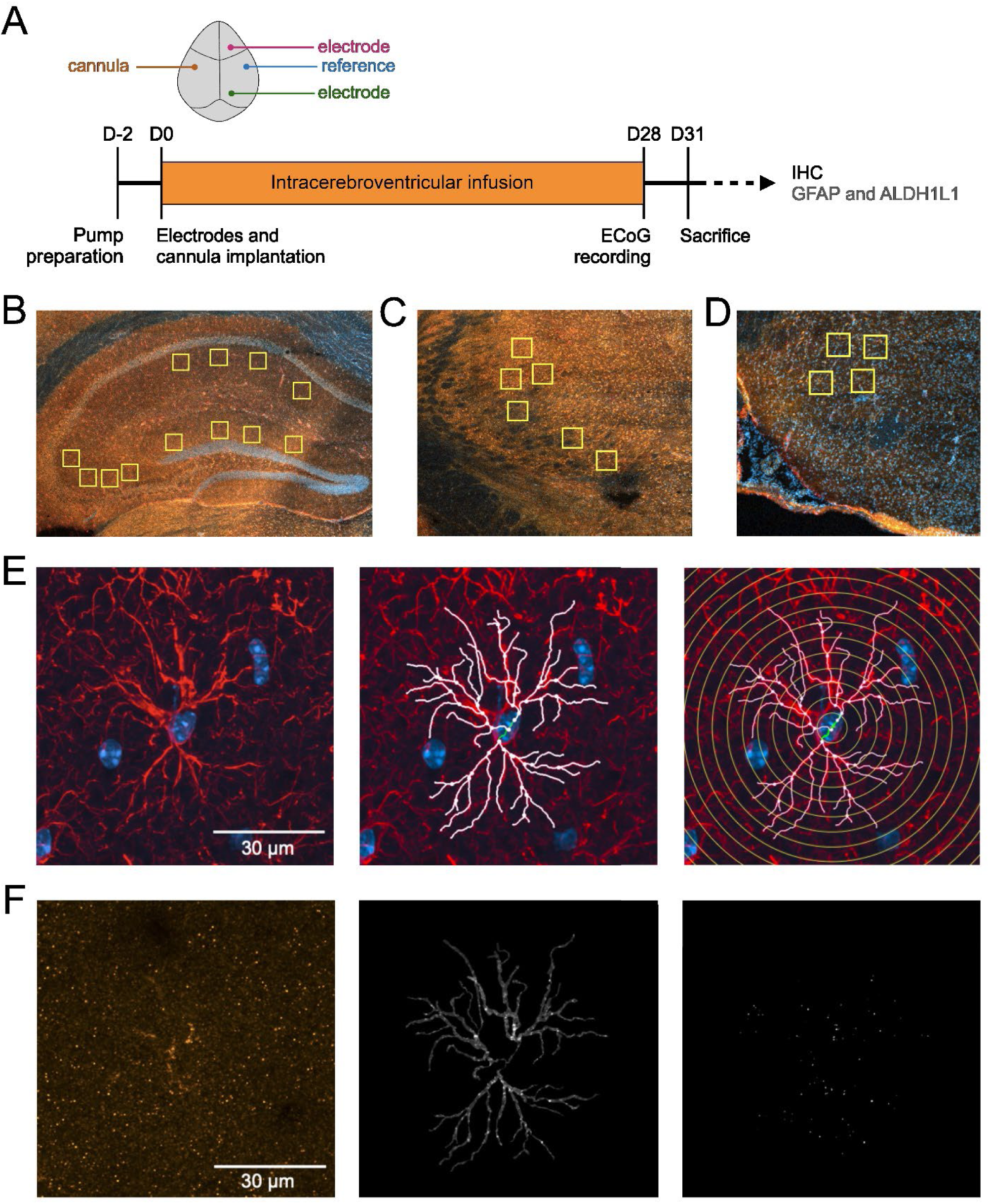
Experimental protocol and methodology related to IHC analyses. (A) Schematic representation of the experimental timeline showing the position of the intracerebroventricular cannula and electrodes, and the duration of SCDi and vehicle treatments. A representative view of the selection of four regions of interest (yellow squares) in hippocampal CA1, CA3 and DG (red staining = GFAP, yellow staining = ALDH1L1, and blue staining = DAPI; also in panels C and D). (B) A representative view of the selection of six regions of interest (yellow squares) in the thalamus (mostly ventral posterolateral and ventral posteromedial nuclei). (C) A representative view of the selection of four regions of interest (yellow squares) in the lateral hypothalamus (LH). (D) A representative image of a GFAP-positive astrocyte (40X objective and 2X zoom, maximum intensity Z-stack projection) on the left, together with the corresponding skeleton of this cell in the middle, and the 4 μm sphere intervals fitted through it for Sholl analysis on the right. (red staining = GFAP, blue staining = DAPI; the 4 μm sphere intervals were centered around the DAPI-stained cell nucleus marked with a green line). (E) A randomly selected frame from ALDH1L1 staining (yellow) Z-stack of the same astrocyte as in (E) is shown on the left image. The middle image shows ALDH1L1 puncta isolated for this frame by applying a fill-in mask of the skeleton shown in panel E. The right image depicts the Gaussian subtraction outcome used for counting ALDH1L1 puncta associated with the corresponding GFAP-positive astrocyte.

### Surgical procedures and ECoG recording

Similar to conducted previously (Hamilton et al., 2015, 2022), osmotic minipumps (Alzet® model 1004; 0.11 μl/hour infusion rate) were filled 48 hours prior to surgical implantation with the SCDi (Abcam #ab142089) dissolved in dimethylsulfoxide (DMSO; Sigma-Aldrich) and diluted in sterile artificial cerebrospinal fluid (aCSF; Harvard Apparatus # 59-7316) for a final infusion concentration of 80 µM. Control/vehicle pumps were filled with the same volume (i.e., 100 µL) of DMSO/aCSF. Osmotic pumps were assembled to a cannula (Plastics One #3280P) via PE tubing immediately after filling. For implantation, mice were deeply anesthetized with ketamine/xylazine (120/10 mg/kg, intraperitoneal injection) combined to continuous isoflurane (0.5-2%, inhalation), and were installed in a stereotaxic frame. Mice were implanted with ECoG and EMG electrodes similar to previously described (El Helou et al., 2013; Areal et al., 2020; Ballester Roig et al., 2023). ECoG electrodes (J.I. Morris Inc. screws #FF00CE125) targeted the motor cortex (1.5 mm anterior to the bregma, 1.5 mm lateral right to the midline) and the visual cortex (1.0 mm anterior to the lambda, 1.5 mm right to the midline), a reference electrode was located above the somatosensory cortex (0.7 mm posterior to the bregma, 2.6 mm right to the midline; Fig. 1A), and EMG electrodes (Delta Scientific 99.99% gold wire 0.2 mm diameter annealed) were implanted in neck muscles. During the same surgery, mice were implanted with the osmotic pump under the skin of their back, and with the cannula for intracerebroventricular targeting with the following coordinates: 0.8 mm posterior to the bregma, 2.3 mm left to the midline, and 2.05 mm below the skull surface with a 27.5° angle left from the vertical plane to avoid interference with ECoG/EMG electrodes. The cannula and electrodes were then fixed to the skull using dental cement (3M RelyX Unicem 2 automix), and electrodes were soldered to a 6-channel connector (ENA AG #BPHF2-O6S-E-3.2). Implanted mice were monitored two times per day for the first three days post-surgery. Just before the end of the 28-day treatment, mice were habituated to cabling conditions for five consecutive days, following which ECoG/EMG signals were recorded under undisturbed conditions at 256 Hz using the Stellate Harmonie software (Natus) and Lamont amplifiers.

### ECoG analyses

Vigilance states (i.e., wakefulness, SWS, PS) were visually assigned to 4-second epochs using bipolar ECoG (motor - visual cortex) and EMG signals as previously described (Ballester Roig et al., 2023; Hector et al., 2023). Artifacts were simultaneously identified and excluded from spectral and scale-free ECoG analyses. The time spent in each vigilance state was averaged for the full 24-hour recording, for the 12-hour light and dark periods, as well as per hour. The number of bouts of different durations was computed for the full 24-hour recording separately for each vigilance state. The bipolar and unipolar (motor and visual cortex separately) ECoG signals were submitted to spectral analysis using a Fast Fourier transform to compute spectral power between 0.5 and 50 Hz with a 0.25 Hz resolution for the full 24-hour recording and/or per 12-hour light and dark periods for each state. Power spectra were expressed as a percent of the mean power of all 0.25-Hz bins of all states for each mouse to interrogate the relative contribution of all frequencies (i.e., relative power spectra). Absolute and relative activity for the full 24-hour recording and 12-hour periods were also calculated for six different frequency bands (i.e., SWA 0.5-5 Hz, Theta 5-9 Hz, Alpha 9-12 Hz, Sigma 12-16 Hz, Beta 16-30 Hz, Gamma 30-50 Hz). In addition, the 24-hour time course of SWA and Theta activity during wakefulness and SWS was calculated using averages per time intervals to which an equal number of epochs contributed and expressed relative to the 24-hour mean activity for each band and each mouse as described previously (Areal et al., 2020; Ballester Roig et al., 2023; Hector et al., 2023).

Parameters related to scale-free ECoG activity were computed using Wavelet-Leaders similar to previously described (Lina et al., 2019), which was developed as a function of earlier methodologies (Ciuciu et al., 2008; Wendt et al., 2009). Briefly, the 1/f^γ^ power decay of the spectral power as a function of frequency plotted in log-log scales was used to compute the scaling exponent γ indicative of the slope of this decay (i.e., γ = 2H + 1) and the Hurst exponent (H). Since the ECoG power spectrum is not monofractal, but rather composed of multiple Hs (Ma et al., 2005; Weiss et al., 2009), the most prevalent Hurst exponent (Hm) and the dispersion (D) of Hs around Hm were extracted using the multifractal Wavelet-Leaders approach. Hm reflects the variability in the long-range dependencies of different ECoG time scales with 0.5 generally considered as the value separating persistence (> 0.5) and anti-persistence (< 0.5). It has been calculated for the artifact-free bipolar signal of each vigilance state for the full 24-hour recording and for the same time intervals as frequency bands described above. The same was done for the multifractal parameter D, with higher absolute D linked to higher complexity (i.e., higher instability and dispersion of Hurst exponents). These two parameters were also computed for the unipolar ECoG signals of each vigilance states for the full 24-hour recording. Two animals had to be excluded from both spectral and scale-free ECoG analyses of the bipolar and visual cortex signals because of predominant artifacts (one WT mouse receiving vehicle and one receiving SCDi). For specific sets of analyses, additional exclusions occurred mainly because of an absence of artifact-free epochs in some intervals (in particular for PS) or of outlier values.

### Immunohistochemistry (IHC)

Twenty-three right hemispheres (n = 6 per experimental group, except n = 5 for 3xTg mice receiving SCDi) were used for IHC to count the number of GFAP- and ALDH1L1-positive cells and conduct a morphological analysis. Thirty µm coronal slices of hemispheres were cut at −18 ± 1°C using a cryostat (Leica #CM3050 S), and slices were directly mounted on Superfrost Plus™ slides (Fisher Scientific #1255015) and stored at −80°C until processing. For IHC, one slice per animal located between 1.94 mm and 2.18 mm posterior to bregma was used to capture a similar region of the thalamus, LH, and dorsal hippocampus. Slides were first placed at −20°C for 8 minutes, and then at room temperature for 4 minutes, following which they were fixed for 30 minutes in 10 % formalin (Sigma-Aldrich #R04586-76), and washed 3 times 5 minutes in phosphate-buffered saline (PBS; Sigma-Aldrich #P5368). Each slide was then dried around the slices, which were isolated using a PAP pen (IHC WORLD #SPM0928). The slides were then incubated for 60 minutes at room temperature in blocking buffer (PBS with 5 % goat serum [Abcam #ab7481] and 0.3 % Triton X-100 [Bio-Rad Laboratories #161-0407]). After removing the blocking buffer and re-applying PAP pen, the slides were incubated overnight at 4°C with primary antibodies (anti-GFAP 1:1000, Invitrogen #PA1-10004; anti-ALDH1L1 1:1000, Abcam ab87117) diluted in PBS with 1 % bovine serum albumin (Winset Inc. #800-095-EG) and 0.3 % Triton X-100. After 4 times 5-minute washes in PBS with 0.5 % Tween® 20 (Fisher Scientific #9005-64-5), each slide was again dried around tissue slices and PAP pen was re-applied. The slides were then incubated 90 minutes in the dark at room temperature with the corresponding secondary antibodies (Alexa Fluor 647 1:1000, Abcam ab150171; Alexa Fluor 555 1:1000, Invitrogen A-21429) diluted in PBS with 1 % bovine serum albumin and 0.3 % Triton X-100. After additional 5-minute washes with 0.5 % Tween 20 in PBS, ProLong® Gold antifade reagent with 4′,6-diamidino-2-phenylindole (DAPI; Invitrogen #P36931) was added to each slide, which were immediately mounted with a glass coverslip (#1.5 thickness; Epredia™ #152455). The slides were sealed with nail polish, and stored in the dark at 4°C until imaging.

### Image acquisition and analyses

All images were acquired using a confocal microscope (Zeiss LSM 900 with Airyscan 2). For cell counting, images were acquired with a 20X lens and a 2X zoom using a Z-stack (average interval between sequential sections of 0.47 µm). Then, non-overlapping regions of interest of the same size were captured: four in the CA1 (stratum radiatum), four in the CA3 (stratum radiatum and lucidum), four in the dentate gyrus (DG; stratum moleculare), six in the ventral posterolateral and ventral posteromedial nuclei of the thalamus, and four in the LH for each animal (Fig. 1B to 1D). Image acquisition was done with a pinhole aperture of one airy unit and the following settings: GFAP excitation = 652 nm, emission = 669 nm; ALDH1L1 excitation = 553 nm, emission = 568 nm; DAPI excitation = 358 nm, emission = 461 nm; collected in all channels. Cell counting was performed for each region of interest on the maximum intensity Z-projection images using the “Cell Counter” plug-in of Fiji (ImageJ 1.54f), and Z-stacks were used for the validation of the counts. A cell was counted when the specific marker (GFAP or ALDH1L1) co-localized with the nuclear stain DAPI. Given our observation of a low GFAP staining in the thalamus and LH, and of a puncta-like ALDH1L1 staining in hippocampal areas, GFAP-positive cells were counted only for hippocampal areas and ALDH1L1-positive cells only in the thalamus and LH. Cell counts were normalized to the size of the counted regions to be reported as cell densities.

A morphological analysis using a skeletonization pipeline applied to the GFAP staining was adapted from a previously described method (Tavares et al., 2017). For this analysis, images were acquired using the same microscope and a 40X oil immersion lens (with a 2X zoom) also with a Z-stack approach (average interval between sequential sections 0.25 µm). A laser power correction was applied through the Z-stack to optimize detail and avoid photobleaching. A total of six astrocytes (non-overlapping regions of size 79.86 x 79.86 µm) per mouse were collected from the CA1 stratum radiatum, and GFAP and DAPI channels were merged. Only astrocytes with clear GFAP-positive processes extending in more than one direction and a DAPI-positive nucleus were considered and skeletonized (Fig. 1E, left panel). The skeletons of the GFAP-positive processes were extracted using the “Simple Neurite Tracer” plug-in of Fiji (Fig. 1E, middle panel), and analyzed using the built-in feature “Measure” to obtain three parameters: radius from the soma (radius in µm from the cell body center to the outermost sphere containing all processes/branches of the astrocyte; see also Sholl analysis below), mean length of processes/branches (µm), and total length of processes/branches (µm). In addition, a Sholl analysis was performed using a sphere radius increment of 4 µm (Fig. 1E, right panel), similar to previously described for astrocytes (Klein et al., 2020; Tavares et al., 2017).

Given the observation of puncta-like ALDH1L1 staining in the CA1 region of the hippocampus, we decided to count the number of ALDH1L1 puncta associated to GFAP-positive astrocytic processes. To do so, the GFAP/DAPI Z-stack was used to define a frame range covering most of the astrocyte, and three frames of the ALDH1L1 Z-stack were randomly chosen within this range (Fig. 1F, left panel). The skeleton of the corresponding astrocyte was then converted into a “mask” using the “Fill out” setting of “Simple Neurite Tracer” at a threshold of 0.005 (threshold enough to cover different branch thicknesses while remaining conservative; arbitrary unit). This mask was then applied to each of the three ALDH1L1 frames to isolate the ALDH1L1 puncta associated with the corresponding astrocyte (Fig. 1F, middle panel). This was done using Gaussian background subtraction with a relative threshold of 0.03 to 0.1 % (Fig. 1F, right panel). ALDH1L1 puncta were then automatically counted, and counts obtained from the three frames were averaged per astrocyte (n = 6 astrocytes per mouse). This method was validated on small segments of the image for which manually counts were compared to the software-generated counts.

### Statistical analyses

Statistical analyses were performed using the software Prism (GraphPad) or Statistica (StatSoft). Two-way analyses of variance (ANOVA) were used with factors Genotype and Treatment to compare wake/sleep variables average per 24-hour recording or 12-hour periods between experimental groups, as well as GFAP- and ALDH1L1-positive cell densities and GFAP-derived morphological parameters. Three-way repeated-measure ANOVA with factors Genotype, Treatment, and light/dark Period, Hour, Interval or Distance from the soma were used to compare the time course of time spent in vigilance states, of activity in frequency bands, of scale-free parameters, and the Sholl analysis-extracted morphology of astrocytes. Tukey’s post hoc tests or planned comparisons were used to decompose significant effects where appropriate, and significance levels of repeated-measure designs were corrected using the Huynh-Feldt correction when needed. Results are presented as mean ± standard error of the mean (SEM), and the threshold for statistical significance was defined as p < 0.05. The size of analyzed samples is indicated in the legend of each figure.

## RESULTS

### Wake/sleep architecture

We first observed that wake/sleep architecture alterations in 3xTg-AD female mice in comparison to WT mice were not significantly impacted by the inhibition of SCD. Mutant mice were found to spend significantly less time awake and more time in SWS during the full 24-hour recording (main Genotype effects wake F1,41 = 13.1, p < 0.001; SWS F1,41 = 14.2, p < 0.001; Fig. 2A), which was driven by specific differences during the dark period (Genotype by Light/Dark Period interaction wake F1,41 = 6.4, p < 0.05; SWS F1,41 = 7.5, p < 0.01; Fig. 2B). SCDi treatment did not significantly impact time spent awake and in SWS (Genotype by Treatment interactions F1,41 < 0.1, p > 0.8; main Treatment effects F1,41 < 1.8, p ≥ 0.2 for the full 24-hour recording), and also did not affect time spent in PS (Genotype by Treatment interaction F1,41 = 0.5, p = 0.5; main Treatment effect F1,41 = 0.1, p = 0.8 for the full 24-hour recording). The 24-hour distribution of wake/sleep states was significantly modified in 3xTg-AD mice, which spent less time awake in the middle of the dark period, more time in SWS during the same hours, and less time in PS around the light to dark transition (Genotype by Hour interactions wake F23,943 = 2.3, p < 0.01; SWS F23,943 = 2.3, p < 0.01; PS F23,943 = 1.6, p < 0.05; Fig. 2C). State distribution across 24 hours was also not significantly changed by SCDi treatment (Genotype by Treatment by Hour interactions F23,943 < 0.9, p > 0.7).

**Fig. 2.**
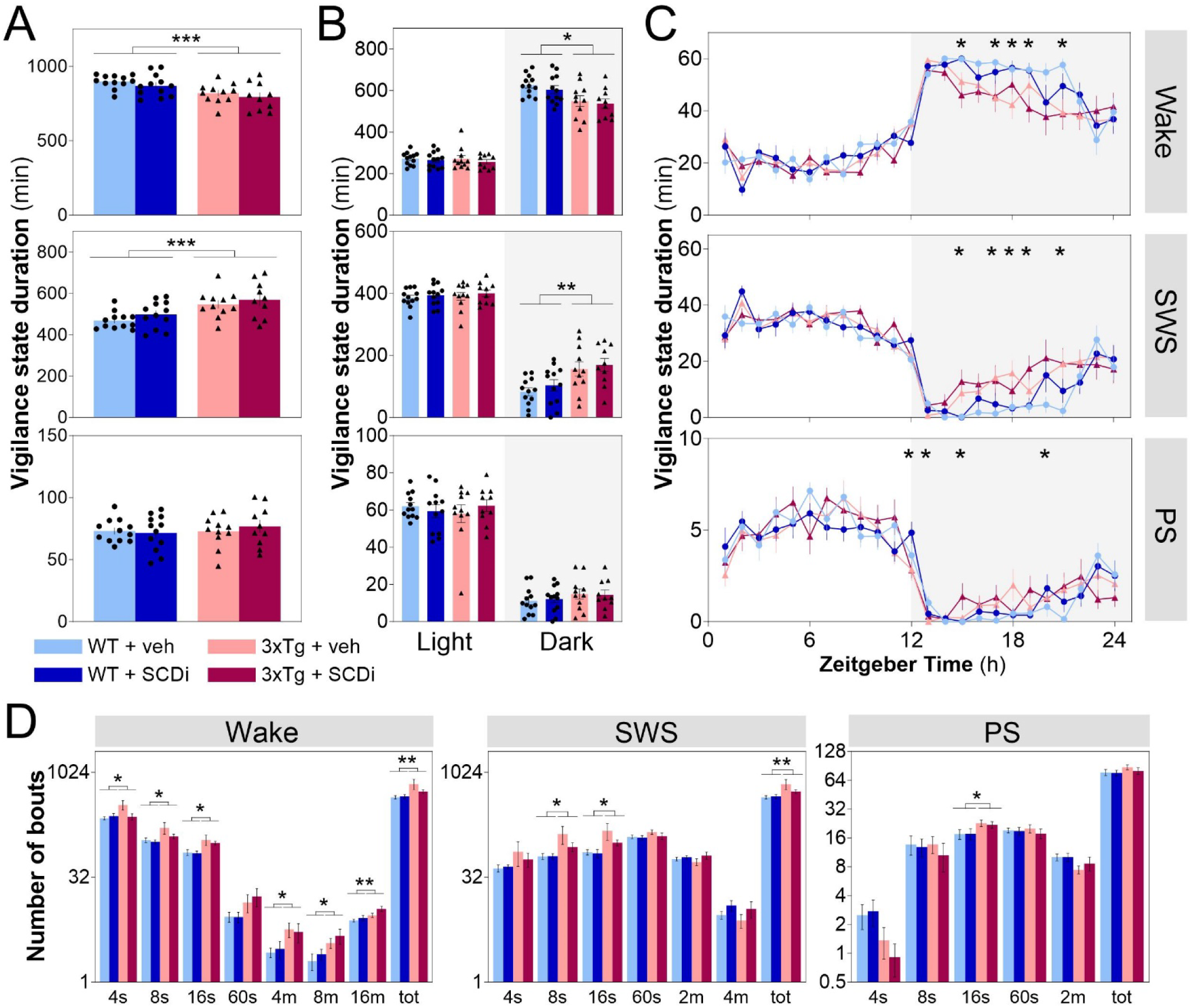
Wake/sleep architecture variables in 3xTg-AD and WT mice treated with SCDi or vehicle. (A) Time spent in wakefulness, SWS and PS during the full 24-hour recording (WT + veh n = 12 mice, WT + SCDi n = 12 mice, 3xTg + veh n = 11 mice, and 3xTg + SCDi n = 10 mice; also in all other panels). ***: p < 0.001 between genotypes. (B) Time spent in wakefulness, SWS and PS during the 12-hour light and the 12-hour dark periods. Grey backgrounds indicate the 12-hour dark period (also in panel C). *: p < 0.05 and **: p < 0.01 between indicated genotypes (also in panels C and D). (C) 24-hour distribution of time spent in wakefulness, SWS and PS. (D) Number of individual bouts of different durations for wakefulness, SWS and PS, and total number of bouts of each state (tot).

In general, mutant mice showed more individual bouts of wake and SWS states (main Genotype effects for total number of bouts: wake F1,41 = 6.9, p = 0.01; SWS F1,41 = 6.8, p = 0.01; Fig. 2D), which is indicative of a higher fragmentation (less consolidation) of vigilance states. This phenotype was also not significantly modified by inhibiting SCD during 28 days (although a trend was observed for very short wake bouts [4-second bouts] to be rescued to WT vehicle values in 3xTg treated with SCDi; Genotype by Treatment interaction F1,41 = 3.5, p = 0.07). These findings reveal major alterations in wake/sleep amount and alternation in 3xTg female mice, including more SWS during the active (dark) period and elevated wake/SWS fragmentation, which are not significantly reverted by the inhibition of SCD during four weeks.

### Spectral analysis of the bipolar ECoG during wake/sleep states

The spectral profile of the ECoG within the different vigilance states, interrogated using a standard spectral analysis, can be considered an indication of state quality. It was initially investigated using the standard bipolar signal for both absolute and relative power because the first informs on, for instance, changes in the organization of the cerebral cortex, whereas the second is indicative of the relative contribution of the different frequency components of the signal. The 24-hour mean absolute power spectra computed between 0.5 and 50 Hz showed significant decreases in spectral power for a wide range of frequencies for the three vigilance states in 3xTg mice in comparison to WT mice (main Genotype effects wake F1,40 > 4.1, SWS F1,40 > 4.2, PS F1,40 > 4.1, all p < 0.05; Fig. 3A). More precisely, when assessing spectral activity in distinct frequency bands, a significant lower activity in 3xTg was found for SWA, Theta, Alpha, Sigma and Beta measured during both wakefulness and SWS (main Genotype effects wake F1,40 > 4.8, p < 0.05; SWS F1,40 > 13.1, p < 0.001; Fig. 3B). During PS, lower activity in mutant mice was significant for Sigma, Beta and Gamma frequency bands (main Genotype effects F1,40 > 6.0, p < 0.05). The absolute activity measured for the six frequency bands was not significantly impacted by SCDi treatment (Genotype by Treatment interactions F1,40 < 2.0, p > 0.1; main Treatment effects F1,40 < 0.9, p > 0.3). These observations indicate a global reduction in ECoG spectral power in 3xTg mice that encompasses all vigilance states.

**Fig. 3.**
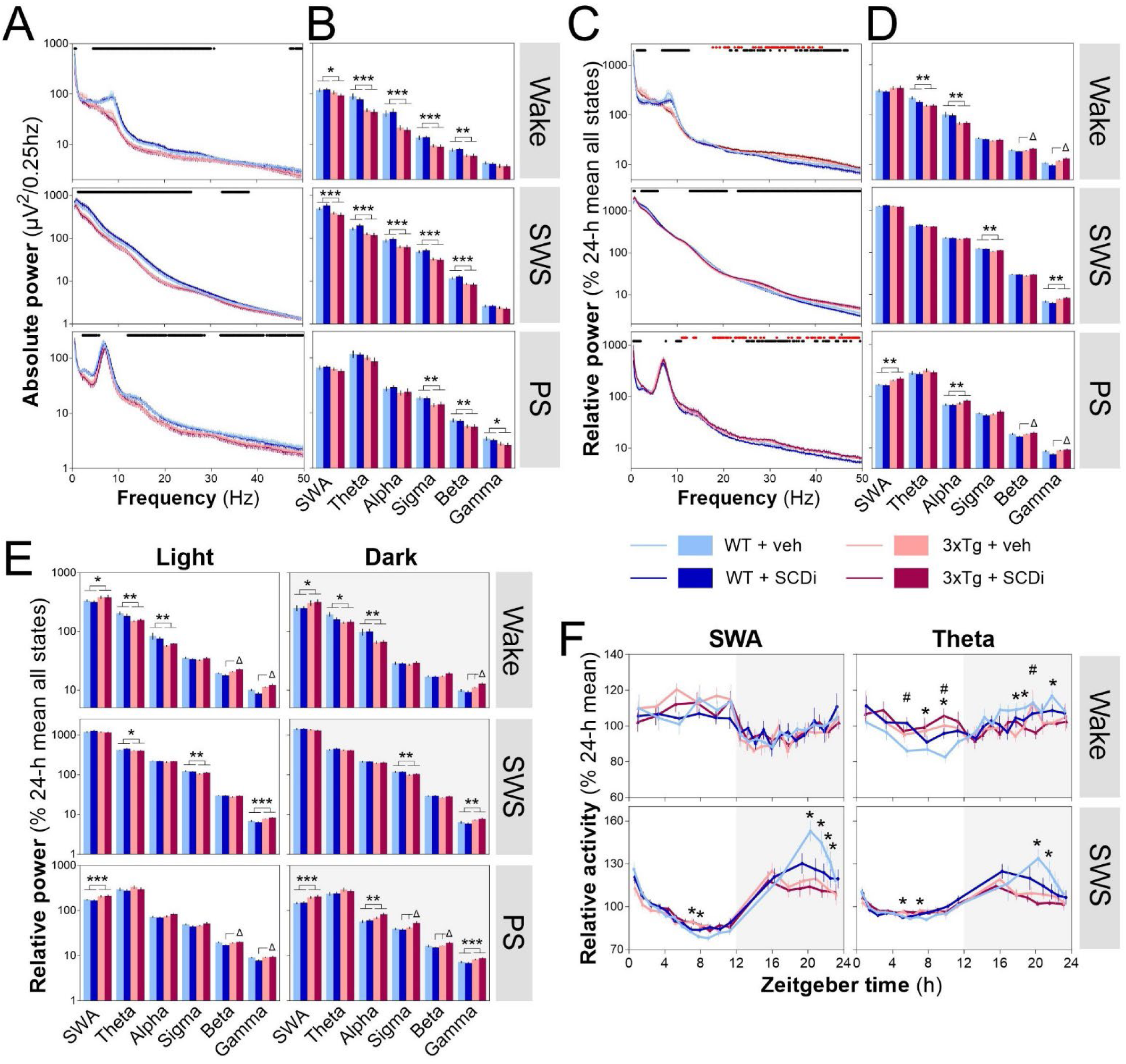
Bipolar ECoG signal during the three vigilance states in 3xTg-AD and WT mice treated with SCDi or vehicle. (A) Mean 24-h power spectra (absolute power) for wake (top), SWS (middle) and PS (bottom). (WT + veh n = 11 mice, WT + SCDi n = 11 mice, 3xTg + veh n = 11 mice, and 3xTg + SCDi n = 10 mice) Black lines on top of graphing area indicate 0.25 Hz-bins with significant differences between genotypes (p < 0.05). (B) Mean 24-h absolute spectral activity for six different frequency bands (SWA 0.5-5 Hz, Theta 5-9 Hz, Alpha 9-12 Hz, Sigma 12-16 Hz, Beta 16-30 Hz, Gamma 30-50 Hz) for wake, SWS and PS (WT + veh n = 12 mice, WT + SCDi n = 11 mice, 3xTg + veh n = 11 mice, and 3xTg + SCDi n = 10 mice; also in panels C to E). *: p < 0.05, **: p < 0.01 and ***: p < 0.001 between genotypes (also for other panels). (C) Mean 24-h power spectra (relative power) for wake (top), SWS (middle) and PS (bottom). Black dots on top of graphing area indicate 0.25 Hz-bin with significant differences between genotypes; red dots bins with significant Genotype by Treatment interactions; and grey dot the bin with a significant difference between treatments (p < 0.05). (D) Mean 24-h relative activity for the six different frequency bands for wake, SWS and PS. Δ: p < 0.05 between indicated groups for bands with significant Genotype by Treatment interactions (also in panel E). (E) Mean 12-h light and 12-h dark relative activity for the six different frequency bands for wake, SWS and PS. Grey backgrounds indicate the 12-hour dark period (also in panel F). (F) 24-h time course of SWA (first column) and theta activity (second column) for wake and SWS (WT + veh n = 11 or 12 mice, WT + SCDi n = 9 or 11 mice, 3xTg + veh n = 10 or 11 mice, and 3xTg + SCDi n = 10 mice; some mice excluded because of no artifact-free epoch for one or two intervals). #: p < 0.05 between SCDi and vehicle.

Interestingly, the 24-hour mean relative power spectra revealed significant effects of SCD inhibition for wakefulness and PS in particular, together with genotype-driven changes in the relative contribution of frequencies during all three states (Genotype by Treatment interactions wake and PS F1,40 4.1, p < 0.05; main Treatment effect PS F1,40 = 6.0, p < 0.02; main Genotype effects all states F1,40 > 4.1, p < 0.05; Fig. 3C). For wakefulness and PS, SCDi treatment impacted the relative contribution of frequencies in the Beta and Gamma bands in a genotype-dependent manner (significant Genotype by Treatment interactions wake F1,40 > 5.1, p < 0.03; PS F1,40 > 4.7, p < 0.04; Fig. 3D), with treated 3xTg mice showing higher relative activity than treated WT mice (p < 0.01). The same observations were made when examining specifically the 12-hour mean relative power of the light period (Genotype by Treatment interactions wake F1,40 > 5.2, p < 0.03; PS F1,40 > 4.9, p < 0.04; Fig. 3E), whereas SCD inhibition affected in a genotype-dependent manner the Gamma band during wakefulness and the Sigma and Beta bands during PS when considering the 12-hour mean relative power of the dark period (Genotype by Treatment interactions wake F1,40 = 5.2, p < 0.03; PS F1,40 > 4.1, p < 0.05; Fig. 3E). These findings suggest that SCDi affects the relative activity in faster frequencies during wake and PS in an opposite manner in 3xTg mice in comparison to WT mice; increasing it in the former and decreasing it in the second.

Concerning genotype differences in the 24-hour mean relative power in the different frequency bands, a significant lower relative contribution in 3xTg mice in comparison to WT mice was found for the Theta and Alpha bands during wakefulness and the Sigma band during SWS (main Genotype effects wake F1,40 > 10.5, p < 0.01; SWS F1,40 = 10.4, p < 0.01; Fig. 3D), whereas the opposite was found for SWS Gamma as well as SWA and Alpha during PS (main Genotype effects SWS Gamma F1,40 = 16.7, p < 0.001; PS F1,40 > 5.6, p < 0.03; Fig. 3D). These differences between genotypes were also generally found when assessing relative power separately for the light and dark periods (main Genotype effects wake F1,40 7.2, p < 0.02; SWS F1,40 > 8.9, p < 0.01; PS F1,40 > 10.9, p < 0.01; Fig. 3E), in addition to more SWA during wake and less Theta during SWS in the light period in 3xTg mice (main Genotype effects wake F1,40 > 4.3, p < 0.05; SWS F1,40 = 4.1, p < 0.05). In sum, mutant mice present major alterations in the relative contribution of different frequency bands during all three vigilance states, alterations that are not significantly rescued by the inhibition of SCD.

The changes in SWA during SWS and Theta activity during wakefulness occurring as a function of the nychthemeron can be considered as index of the homeostatic process that regulates sleep (Vassalli and Franken, 2017). The 24-hour variations in SWA and Theta activity measured during wakefulness and SWS were thus compared between groups to investigate the impact on the daily dynamics of the activity in these frequency bands (Fig. 3F). The dynamics of SWA during SWS and of Theta activity during wake and SWS was significantly impacted by genotype (Genotype by Interval interactions wake F17,629 = 2.6, p < 0.05; SWS F17,663 > 2.9, p < 0.05), with higher activity near the middle of the light period and lower activity during the dark period in 3xTg mice in comparison to WT mice, resulting in a generally lower amplitude of daily variations in mutant animals. SCDi treatment also decreased the daily variations in Theta activity during wakefulness (Treatment by Interval interaction F17,629 = 1.7, p < 0.05), by way of increasing activity during the mid-light period and decreasing it in the dark in comparison to vehicle. These findings suggest that genotype and treatment impact 24-hour dynamics of slower ECoG activity in independent manners.

### Spectral analysis of the motor and visual ECoG

We then aimed at verifying whether the inhibition of SCD and the 3xTg model affect ECoG activity in a brain region-specific manner. Group comparisons were thus conducted separately for the motor and visual cortex signals using relative activity expressed as a function of the mean of WT mice receiving vehicle. For the motor cortex, SCDi treatment significantly impacted activity during wakefulness in a genotype-dependent manner for few isolated Hz-bins during the light period and for frequencies below 18 Hz during the dark period (Genotype by Treatment interactions F1,38 > 4.1, p < 0.05; Fig. 4A), with treatment altering activity in only one genotype or in opposite manner in 3xTg and WT mice. SCD inhibition also decreased activity during PS for several Hz-bins around 20 Hz and above 40 Hz during both the light and dark periods (main Treatment effects F1,38 > 4.1, p < 0.05; Fig. 4A). Motor cortex activity was importantly affected by genotype in all states and 12-hour periods with, in general, mutant mice showing less activity below 20 Hz and higher activity above 20 Hz in comparison to WT mice, except for PS for which activity between 7 and 10 Hz was significantly higher in mutant mice (main Genotype effects wake F1,38 > 4.2, SWS F1,38 > 4.1, PS F1,38 > 4.1, all p < 0.05; Fig. 4A). For the visual cortex, SCD inhibition generally decreased activity below 20 Hz during wake in WT mice but increased it (or did not change it) in 3xTg mice (Genotype by Treatment interactions F1,36 > 4.1, p < 0.05; Fig. 4B). However, the opposite was found for activity below 6 Hz during SWS, with SCDi decreasing it in 3xTg mice and increasing in WT mice (Genotype by Treatment interactions F1,36 > 4.2, p < 0.05; Fig. 4B). These findings apply to both the light and dark periods. In addition, SCD inhibition was found to significantly decrease activity during PS for multiple Hz-bins above 25 Hz for the light and dark periods (main Treatment effects F1,36 > 4.1, p < 0.05; Fig. 4B). Mutant mice also showed multiple alterations of activity for the visual cortex signal measured during the light and dark periods. During wakefulness, 3xTg mice generally expressed higher activity in lower (< 6 Hz) and higher (> 20 Hz) frequencies but less between 7 and 10 Hz in comparison to WT mice (main Genotype effects F1,36 > 4.1, p < 0.05; Fig. 4B). During SWS, 3xTg mice showed significantly less activity than WT mice below 22 Hz, but more above 26 Hz (main Genotype effects F1,36 > 4.2, p < 0.05; Fig. 4B). Finally, 3xTg mice also had in general more activity than WT mice in multiple Hz-bins between 0.5 and 50 Hz in PS (main Genotype effects F1,36 > 4.1, p < 0.05; Fig. 4B). These findings indicate that the inhibition of SCD alters synchronized activity in a genotype-, frequency-, state- and topographic site-dependent manner, and that mutant mice show a highly specific spectral signature that also depends on frequency, state and brain area.

**Fig. 4.**
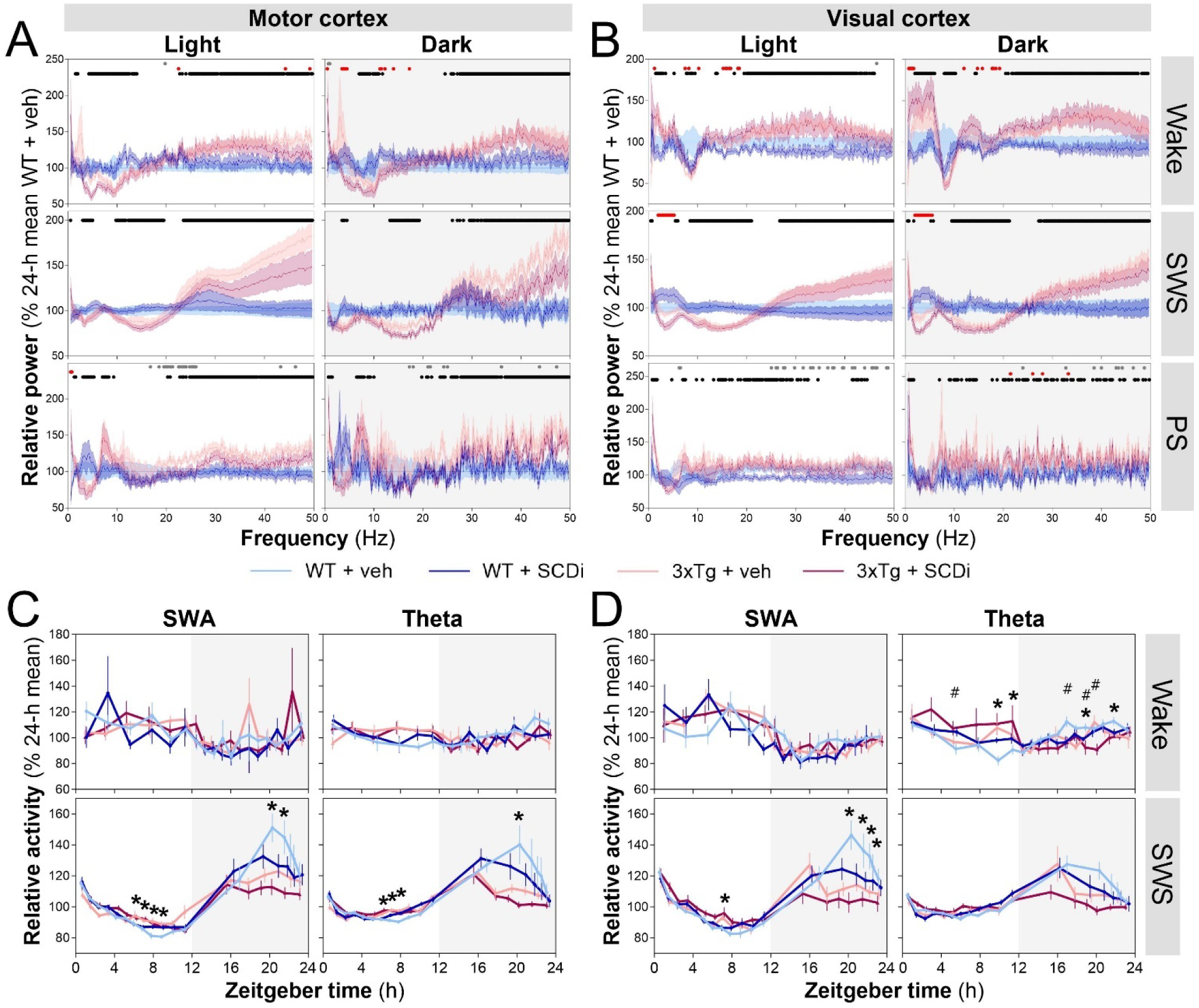
Motor and visual cortices ECoG signals during vigilance states in 3xTg-AD and WT mice treated with SCDi or vehicle. (A) Mean 12-h spectral power for the motor cortex represented as a percent of WT + vehicle mice separately for the light and dark periods for wakefulness (top), SWS (middle) and PS (bottom). Grey backgrounds indicate the 12-hour dark period (also in other panels). (WT + veh n = 12 mice, WT + SCDi n = 10 or 11 mice, 3xTg + veh n = 10 mice, and 3xTg + SCDi n = 8 or 9 mice; some animals excluded because of outlier values) Black dots on top of graphing area indicate 0.25 Hz-bins with significant differences between genotypes; red dots bins with significant Genotype by Treatment interaction; and grey dots the bins with significant differences between treatments (p < 0.05; also in panel B). (B) Mean 12-h spectral power for the visual cortex represented as a percent of WT + vehicle mice separately for the light and dark periods for wake, SWS and PS. (WT + veh n = 11 mice, WT + SCDi n = 11 mice, 3xTg + veh n = 10 mice, and 3xTg + SCDi n = 7 to 9 mice; some animals excluded because of outlier values) (C) Motor cortex 24-h time course of SWA (0.5-5 Hz; first column) and theta activity (5-9 Hz; second column) for wake (top) and SWS (bottom). (WT + veh n = 10 or 11 mice, WT + SCDi n = 8 to 10 mice, 3xTg + veh n = 10 mice, and 3xTg + SCDi n = 8 or 9 mice; some animals excluded because of no artifact-free epoch for some intervals or because of outlier values) *: p < 0.05 between genotypes (also in panel D). (D) Visual cortex 24-h time course of SWA (first column) and theta activity (second column) for wake and SWS (WT + veh n = 10 or 11 mice, WT + SCDi n = 9 or 11 mice, 3xTg + veh n = 10 mice, and 3xTg + SCDi n = 7 or 8 mice; some animals excluded because of no artifact-free epoch for some intervals or because of outlier values). #: p < 0.05 between SCDi and vehicle.

When examining the 24-hour dynamics of SWA and Theta activity during wake and SWS at the level of the motor and visual cortex, significant differences between treatments and genotypes were also found. More precisely, the time course of SWS SWA and Theta activity was altered in 3xTg in comparison to WT mice for the motor cortex (Genotype by Interval interactions F17,595 > 4.5, p < 0.02; Fig. 4C), with a lower amplitude of day-night changes in mutant mice. A similar observation was made for SWS SWA 24-hour dynamics of the visual cortex (Genotype by Interval interaction F17,595 = 5.7, p < 0.001; Fig. 4D), whereas it did not reach statistical significance for SWS Theta activity of the visual cortex (Genotype by Interval interaction F17,595 = 1.7, p > 0.1; Fig. 4D). Concerning wakefulness, only the time course of visual cortex Theta activity was impacted by genotype and treatment, showing that 3xTg mice have more activity before transition to dark and less activity during the second half of the dark period in comparison to WT mice (Genotype by Interval interaction F17,561 = 2.5, p = 0.01), and SCDi increasing activity during the mid-light but decreasing it during the mid-dark in comparison to vehicle (Treatment by Interval interaction F17,561 = 2.3, p = 0.02). These findings indicate that genotype impacts the 24-hour dynamics of slower ECoG activity for both the motor and visual cortex, and that treatment modifies the dynamics of wake Theta activity for the visual cortex only.

### Scale-free activity of the wake/sleep ECoG

The scale-free activity of the ECoG can further inform on signal quality (i.e., signal dynamics complexity) of the different states by assessing the relationship between activity at different time scales. As described in the methodology, two parameters have been extracted from the ECoG bipolar and unipolar signals, namely Hm, indicative of the persistence across time scales, and D, representing the dispersion around Hm and the degree of multifractality. For the bipolar signal, the 24-hour mean Hm was significantly lower in 3xTg than in WT mice for wakefulness and SWS, but not for PS (main Genotype effects wake F1,39 = 23.1, p < 0.001; SWS F1,39 = 47.7, p < 0.001, PS F1,39 = 1.2, p > 0.05; Fig. 5A, first column). The 24-hour dynamics of Hm was also found to differ between genotypes for wakefulness and SWS, but not for PS (Genotype by Interval interaction wake F17,612 = 3.1, p < 0.001; SWS F17,646 = 44.3, p < 0.001; PS F11,286 = 1.1, p = 0.4; Fig. 5A, second column). More precisely, lower Hm in 3xTg mice in comparison to WT mice appeared more prominent during the dark period for wakefulness, and at the beginning of the light period and end of the dark period for SWS. There was no significant effect of the SCDi treatment on the Hm 24-hour mean or 24-hour dynamics for all three states (24-hour mean: Genotype by Treatment interactions F1,39 < 1.4, p > 0.2, and main Treatment effects F1,39 < 0.7, p > 0.4; 24-hour dynamics: Genotype by Treatment by Interval interactions F11/17,286/612/646 < 0.9, p > 0.6; Genotype by Treatment interactions F1,26/36/38 < 1.6, p > 0.2; and main Treatment effects F1,26/36/38 < 0.9, p 0.3).

**Fig. 5.**
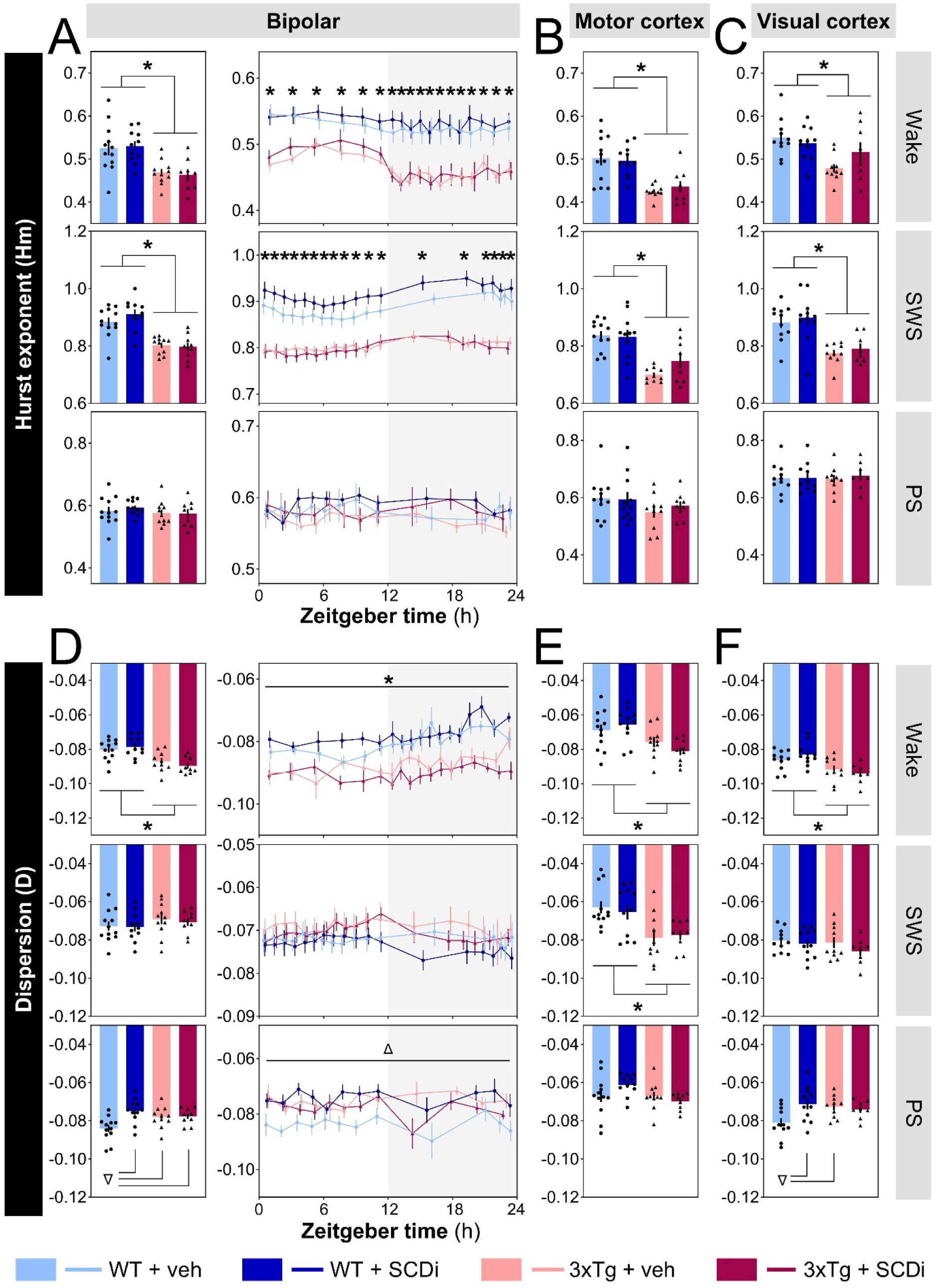
Scale-free activity of the ECoG during wakefulness, SWS and PS in 3xTg-AD and WT mice treated with SCDi or vehicle. (A) Predominant Hurst exponent (Hm) of the bipolar ECoG signal, computed for the full 24-h recording (left), and per time interval (right) for wake (top), SWS (middle) and PS (bottom). Grey backgrounds indicate the 12-hour dark period (also in panel D). (full 24-h recording WT + veh n = 12 mice, WT + SCDi n = 11 mice, 3xTg + veh n = 10 or 11 mice, and 3xTg + SCDi n = 9 mice [also for panels B, D and E]; for the analyses per interval WT + veh n = 7 to 12 mice, WT + SCDi n = 9 to 11 mice, 3xTg + veh n = 6 or 10 mice, and 3xTg + SCDi n = 8 or 9 mice [also for panel D]; some animals excluded because of no artifact-free epoch for some intervals, in particular for PS, or because of outlier values) *: p < 0.05 between genotypes (also in all other panels). (B) Hm computed for the full 24-h recording during wake (top), SWS (middle) and PS (bottom) for the motor cortex. (C) Hm computed for the full 24-h recording during the three states for the visual cortex. (WT + veh n = 11 mice, WT + SCDi n = 11 mice, 3xTg + veh n = 10 mice, and 3xTg + SCDi n = 8 mice; also for panel F). (D) Dispersion (D) of Hurst exponents around Hm for the bipolar ECoG signal, computed for the whole 24-h recording (left), and per time interval (right) for wake (top), SWS (middle) and PS (bottom). Δ: p < 0.05 between indicated groups (also in F) or between WT + veh and other groups. (E) D computed for the full 24-h recording during wake (top), SWS (middle) and PS (bottom) for the motor cortex. (F) D computed for the full 24-h recording during the three states for the visual cortex.

A lower Hm in 3xTg mice suggests more anti-persistence (less persistence) across time scales of the ECoG signal during wakefulness and SWS. Interestingly, the same observation was made when analyzing the 24-hour mean Hm of the motor cortex and visual cortex separately (main Genotype effects motor cortex wake F1,38 = 30.4, p < 0.001 and SWS F1,38 = 34.2, p < 0.001; visual cortex wake F1,36 = 10.3, p < 0.05 and SWS F1,36 = 28.8, p < 0.001; Fig. 5B and C). Once again, no significant difference between genotypes was found for PS (main Genotype effects motor cortex F1,38 = 2.7, p = 0.1; visual cortex F1,36 = 0.04, p > 0.8) and no significant effect of treatment for any state (Genotype by Treatment interactions F1,36/38 < 3.2, p > 0.08; and main Treatment effects F1,36/38 < 1.3, p > 0.1). This suggest that the genotype effect on Hm is state-dependent and relatively global, observable for two distant regions of the cerebral cortex.

The 24-hour mean D was also found to differ between genotypes in a state-dependent manner. More precisely, D was found to be significantly more negative (i.e., more multifractal) in 3xTg mice in comparison to WT mice for the bipolar ECoG signal during wakefulness but not during SWS (main Genotype effects wake F1,38 = 25.6, p < 0.001; SWS F1,39 = 1.6, p = 0.2; Fig. 5D, first column). Similar observations were made for the wakefulness D measured at the level of the motor cortex and the visual cortex (main Genotype effects motor F1,38 = 13.7, p < 0.001; visual F1,36 = 16.4, p < 0.001; Fig. 5E and F, first row), whereas SWS D showed a significantly more negative value in 3xTg mice for the motor cortex but no significant genotype effect for the visual cortex (main Genotype effects motor F1,38 = 15.5, p < 0.001; visual F1,36 = 1.2, p = 0.3; Fig. 5E and F second row). The more negative wakefulness D for the bipolar signal in 3xTg mice in comparison to WT mice was independent of time-of-day as indicated by the 24-hour dynamics (main Genotype effect F1,36 = 29.0, p < 0.001; Genotype by Interval interaction F17,612 = 1.1, p = 0.3; Fig. 5D, second column). For wakefulness and SWS, SCDi treatment did not impact D for the three ECoG signals (Genotype by Treatment interactions F1,36/38/39 < 2.1, p > 0.1; and main Treatment effects F1,36/38/39 < 1.7, p > 0.2).

Regarding D measured during PS, the 24-hour mean of the bipolar signal was found to be impacted by SCDi treatment in a genotype-dependent manner. Indeed, WT mice receiving SCDi showed a significantly less negative D in comparison to WT mice receiving vehicle while the treatment did not significantly impact D in 3xTg mice that are already showing a less negative D (Genotype by Treatment interaction F1,39 = 6.1, p < 0.02; Fig. 5D, last row). This effect was also found when considering the visual cortex (Genotype by Treatment interaction F1,36 = 5.5, p < 0.03; Fig. 5F, last row) but not the motor cortex (Genotype by Treatment interaction F1,38 = 3.2, p = 0.08; main Genotype effect F1,38 = 3.3, p = 0.08; main Treatment effect F1,38 = 0.5, p = 0.5; Fig. 5E, last row). There was no effect of time-of-day on D measured during PS (Genotype by Treatment by Interval interaction F11,286 = 1.2, p = 0.3; Genotype by Interval interaction F11,286 = 0.5, p = 0.9; Fig. 5D, last row of second column). Overall, analyses of D reveal state-specific changes, indicative of more fractal dispersion (more multifractality) during wake in particular in 3xTg mice, and less dispersion during PS after SCDi treatment in WT mice.

### Astrocyte-specific markers

The number of cells expressing two well-recognized markers of astrocytes (and of astrocyte activation in the case of GFAP) were investigated after SCD inhibition in WT and 3xTg mice. In the hippocampus, GFAP-positive cell density was found to be modified by both treatment and genotype in a region-dependent way. In CA1, SCDi treatment significantly restored the elevated cell density in 3xTg mice (Genotype by Treatment interaction F1,88 = 4.9, p = 0.03; Fig. 6A). In CA3, GFAP-positive cell density was observed to be higher in 3xTg in comparison to WT mice (main Genotype effect F1,88 = 4.6, p < 0.04; Fig 6B), and SCDi did not significantly impact it (Genotype by Treatment interaction F1,88 < 0.1, p = 0.9; main Treatment effect F1,88 = 1.3, p = 0.3). For the DG, no significant effect of genotype and/or treatment was found on GFAP-positive cell density (Genotype by Treatment interaction F1,88 = 0.4, p = 0.5; main Genotype effect F1,88 = 0.7, p = 0.4; main Treatment effect F1,88 = 0.6, p = 0.4; Fig. 6C). These findings support an elevated number of GFAP-stained cells in the CA1 and CA3 regions of the hippocampus in 3xTg mice that can be rescued to WT values in CA1 by the inhibition of SCD.

**Fig. 6.**
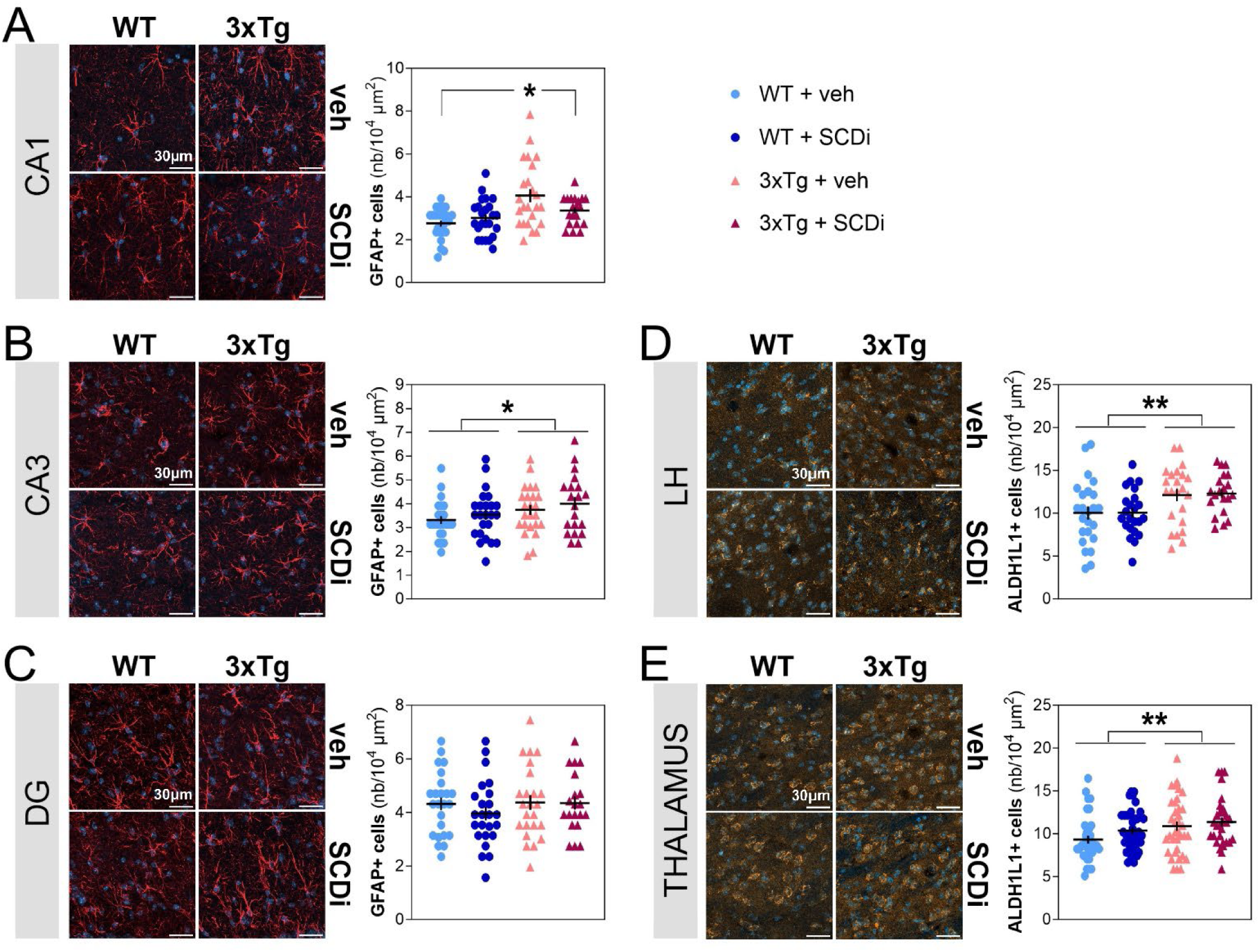
GFAP-positive cell counts in the hippocampus and ALDH1L1-positive cell count in the thalamus and LH in 3xTg-AD and WT mice treated with SCDi or vehicle. (A) Representative images (left) and quantification (right) of GFAP-positive astrocytes in CA1. The red staining is GFAP, and the blue staining DAPI (also in panels B and C). The quantification was computed using the four regions of interest of each animal as different datapoints. *: p < 0.05 between indicated groups. (WT + veh n = 6 mice, WT + SCDi n = 6 mice, 3xTg + veh n = 6 mice, and 3xTg + SCDi n = 5 mice; also in all other panels) (B) Representative images (left) and quantification (right) of GFAP-positive astrocytes in CA3. The quantification was computed using the four regions of interest of each animal as different datapoints. *: p < 0.05 between 3xTg and WT mice. (C) Representative images (left) and quantification (right) of GFAP-positive astrocytes in the dentate gyrus (DG). The quantification was computed using the four regions of interest of each animal as different datapoints. (D) Representative images (left) and quantification (right) of ALDH1L1-positive astrocytes in the LH. The yellow staining is ALDH1L1, and blue staining DAPI (also in panel E). The quantification was computed using the four regions of interest of each animal as different datapoints. **: p < 0.05 between 3xTg and WT mice (also in E). (E) Representative images (left) and quantification (right) of ALDH1L1-positive astrocytes in the thalamus. The quantification was computed using the six regions of interest of each animal as different datapoints.

Concerning the LH and targeted thalamic nuclei, the density of ALDH1L1-positive cells was also significantly elevated in 3xTg in comparison to WT mice (main Genotype effects F1,86/133 > 7.0, p < 0.01; Fig. 6D and 6E). However, SCDi treatment did not affect ALDH1L1-positive cell density in the LH and thalamus (Genotype by Treatment interactions F1,86/133 < 0.4, p > 0.5; main Treatment effects F1,86/133 < 2.7, p > 0.1). The important influence of genotype on ALDH1L1 staining in these sleep-related regulatory brain regions is thus not modulated by 28 days of SCD inhibition.

### GFAP-derived astrocyte morphology

The morphology of CA1 astrocytes was found to be modified by SCD inhibition in a manner independent of genotype. Indeed, the SCDi treatment significantly increased the radius size of the area covered by astrocytes (main Treatment effect F1,134 = 6.0, p = 0.02; Fig. 7A and 7B), and showed a tendency to increase the mean length of astrocytic processes/branches (F1,134 = 3.0, p < 0.09; Fig. 7C); whereas these parameters were not significantly affected by genotype (Genotype by Treatment interactions F1,134 < 0.2, p > 0.7; main Genotype effects F1,134 < 1.3, p > 0.2). However, the total length of astrocytic processes/branches was not significantly modified by treatment and/or genotype (Genotype by Treatment interaction F1,134 = 1.5, p = 0.2; main Treatment effect F1,134 = 0.3, p = 0.6; main Genotype effect F1,134 = 2.1, p = 0.15; Fig. 7D), which also applies to the number of branch intersections as a function of the distance from the soma (interactions with factors Genotype and Treatment F15,285 < 0.5, p>0.9; main Genotype and Treatment effects F1,19 < 0.8, p > 0.3; Fig. 7E). These observations suggest that the inhibition of SCD for 28 days modestly increases the length of individual astrocytic processes resulting in a generally larger diameter of coverage.

**Fig. 7.**
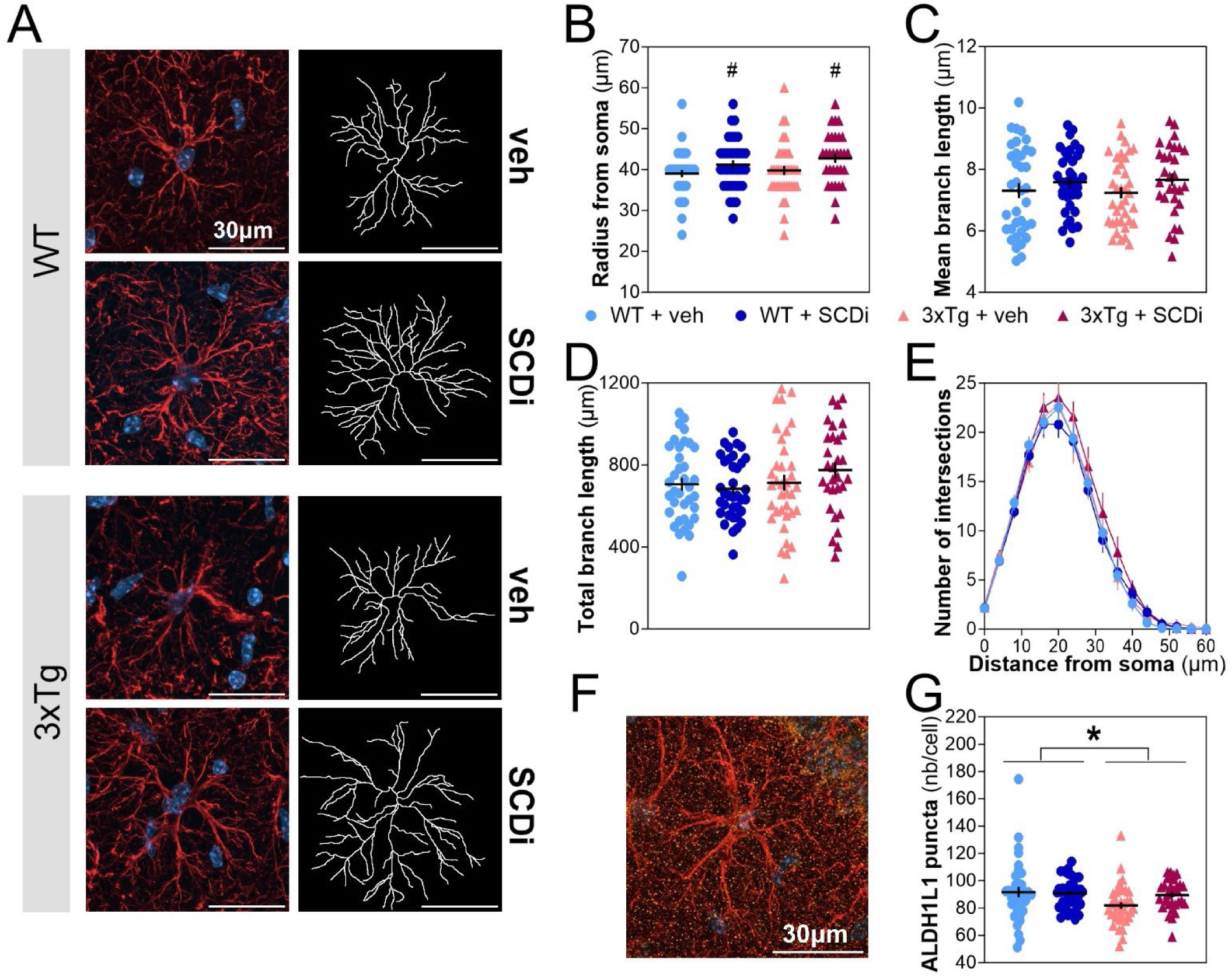
GFAP-derived astrocyte morphology and number of ALDH1L1 puncta per GFAP-positive astrocyte in the CA1 of 3xTg-AD and WT mice treated with SCDi or vehicle. (A) Representative images of GFAP-positive astrocytes (red staining = GFAP, blue staining = DAPI) on the left and corresponding skeletons on the right for one animal of each group. (B) Length of radius shown for six astrocytes per animal for the four experimental groups (WT + veh n = 6 mice, WT + SCDi n = 6 mice, 3xTg + veh n = 6 mice, and 3xTg + SCDi n = 5 mice; also in panels C, D, E and G). #: p < 0.05 between SCDi and vehicle (main Treatment effect). (C) Mean length of astrocytic processes plotted for six astrocytes per animal for the four groups. (D) Total length of astrocytic processes shown for six astrocytes per animal for the four groups. (E) Number of intersections as a function of the distance from the soma averaged for the six astrocytes per animal for the four experimental groups. (F) Representative image of the puncta-like ALDH1L1 staining of a GFAP-positive astrocyte in CA1 for a 3xTg mice treated with SCDi (red staining = GFAP, yellow staining = ALDH1L1, and blue staining = DAPI). (G) Number of ALDH1L1 puncta per astrocyte shown for six astrocytes per mouse for each group. *: p < 0.05 between 3xTg and WT mice (main Genotype effect).

In the CA1 region of the hippocampus, the ALDH1L1 staining mainly appears in the form of small puncta (Fig. 7F). The number of ALDH1L1 puncta associated to GFAP-stained processes/branches and cell bodies was found to be significantly lower in 3xTg mice in comparison to WT mice (main Genotype effect F1,134 = 4.2, p = 0.04; Fig. 7G). The number of puncta per GFAP-stained cell was not significantly impacted by SCDi treatment (Genotype by Treatment interaction F1,134 = 2.5, p = 0.12; main Treatment effect F1,134 = 1.6, p = 0.2). In summary, SCD inhibition impacted astrocyte morphology in CA1, while genotype affected (independently of treatment) ALDH1L1 staining associated to astrocytes in the CA1 hippocampal region.

## DISCUSSION

We have here uncovered that targeting lipid metabolism using the inhibition of SCD impacts the quality of wake/sleep states as assessed using spectral and multifractal analyses of the ECoG, but not the time spent and fragmentation/consolidation of wake/sleep states. More precisely, we found that SCD inhibition changes the ECoG spectral signatures of wakefulness and PS when considering the motor cortex, and spectral signatures of wake, SWS and PS when considering the visual cortex. The effects of this treatment were similar in 3xTg-AD and WT mice during PS, but were generally opposite or different in the two genotypes during wakefulness and SWS, which also applies to multifractal complexity of the PS ECoG signal of the visual cortex that was reduced by SCD inhibition only in WT mice. Globally, these observations indicate that the inhibition of SCD is not noticeably correcting the wide range of modifications in wake/sleep variables found in 3xTg-AD mice. However, we discovered that this lipid metabolism-targeting treatment rescues the elevated number of GFAP-stained cells in hippocampal CA1, without restoring changes observed to astrocyte-specific makers in CA3, LH and thalamic nuclei. The overall findings are suggesting that the selected treatment and the 3xTg-AD model are impacting the quality of wake/sleep states in separated manners, and that the inhibition of SCD shows promise in reverting changes in astrocytic functions.

Our comprehensive investigation of wake/sleep phenotypes has allowed to refine previous observations in 3xTg-AD mice in addition to identifying novel features characterizing this model. Firstly, we found that 5-month old mutant animals sleep more during the dark (active) period, which is in line with findings using a piezoelectric evaluation in 16-month old females (Saber et al., 2021). Our data indicate that this is specifically linked to more time spent in SWS. However, no change in time spent awake and asleep was found in 18-month old 3xTg mice when considering the two sexes (i.e., 4 females and 4 males; Kent et al., 2018), which could indicate a sex-specific effect or an effect that can be captured only at a younger age. More sleep during the active period in 3xTg-AD females is reminiscent of the lower amplitude of the wake/sleep cycle in AD patients comprising a higher incidence of daytime naps (Wang and Holtzman, 2020; Kent et al., 2021), but differs from several other transgenic mouse models that generally show more time spent awake and less time asleep (Dufort-Gervais et al., 2019). Given the reported wake-promoting role of LH astrocytes (Clasadonte et al., 2017; Cai et al., 2022), alterations in astrocytic function in the LH could be a factor contributing to the diminished wake amount in 3xTg-AD mice. Our observation of an elevated number of ALDH1L1-positive cells in the LH could indeed suggest an impaired function of astrocytes (e.g., sign of abnormal compensation or of overactivation) preventing these cells from effectively promoting wake.

Secondly, we exposed multiple alterations in ECoG rhythmic and scale-free activity in the selected AD mouse model. Indeed, absolute power was globally reduced (i.e., all states) and a redistribution of the contribution of activity in different frequencies in favor of a higher contribution of faster frequencies (particularly manifest for SWS) was characterizing adult 3xTg-AD female mice, together with more anti-persistent scale-free activity (wake and SWS). These findings could be specific for the targeted age because an absence of change in ECoG relative activity of the three vigilance states for similar cortical regions was reported in 3xTg-AD mice of both sexes studied at 18 months (Kent et al., 2018). The reduction of absolute power could be considered consistent with the reduced number of sleep oscillations (i.e., spindles and sharp-wave ripples) previously reported for this model after a learning task (Benthem et al., 2020). In addition, more anti-persistent ECoG multifractal activity may indicate a less organized/coordinated neuronal network in 3xTg-AD mice, which is also supported by observations of impaired hippocampal-parietal and delta-spindle coupling after learning (Benthem et al., 2020). Our findings are suggestive of an overall increased scale-free complexity in mutant mice. Given that reports in AD patients using related metrics are most often indicative of a decreased arrhythmic/fractal complexity of the electroencephalogram (Lau et al., 2022; Averna et al., 2023), it is interesting to note that the deviation from the ‘optimal range’ in the complexity of electroencephalographic activity has been proposed to be a more suitable variable to use for clinical applications in comparison to the directionality of the changes (Lau et al., 2022). It is anticipated that a decrease in dendritic spine density in the targeted cerebral cortex regions, as reported for the hippocampus in this model (Hamilton et al., 2022), or other types of synaptic dysfunctions are contributing to the extensive modifications in ECoG rhythmic and arrhythmic activity in 3xTg-AD animals.

Our hypothesis that the inhibition of SCD could rescue wake/sleep alterations in 3xTg-AD mice (via a mechanism implicating astrocytes) was not confirmed by data resulting from wake/sleep phenotyping. SCDi did not impact wake/sleep architecture variables, but did modify ECoG rhythmic activity particularly when considering the motor and visual cortex separately. During PS, SCDi reduced ECoG activity (and multifractal dispersion in WT), whereas it changed wake and SWS rhythmic activity in different manners in 3xTg-AD and WT mice. This implies that this treatment can modulate wake/sleep quality, and in a manner that depends on how the brain is wired and affected by AD-related pathology. The inhibition of SCD could impact ECoG activity via changing synaptic strength since it was found to affect the density of dendritic spines in the hippocampus (Hamilton et al., 2022). In parallel, SCDi could modulate the ECoG by altering fatty acid concentration/composition because fatty acid administration and the activation of a fatty acid receptor were shown to affect neuronal firing (Arsenault et al., 2011; Barki et al., 2022). Despite the very few available studies investigating the direct impact of lipids on wake/sleep quality, it is of interest to observe that SCD effects on ECoG activity were dependent on wake/sleep state, which suggest effects on state-specific regulatory cell populations. More research has rather focused on the impact of sleep disturbances on lipids (Bell et al., 2013; Broussard et al., 2015). In fact, sleep deprivation was shown to increase markers of SCD activity in the serum of healthy young men (Skuladottir et al., 2016). Accordingly, disturbed sleep could contribute to elevated MUFAs in the AD brain, and inhibiting SCD may represent a way to counteract the effect of sleep disturbances in AD.

As presented in the introduction, astrocytes are in a key position to mediate a relationship between wake/sleep states and lipid metabolism. Previous work reported changes in astrocytes in 3xTg-AD mice that depend on brain regions. Indeed, increased GFAP staining (i.e., fluorescence intensity) and other indications of astrocytic hyperreactivity have been reported in primary cell cultures from the 3xTg-AD cerebral cortex (González-Molina et al., 2021), while atrophic astrocytes were found in the entorhinal cortex of 3xTg mice starting at 1 month of age (Yeh et al., 2011), and signs of atrophy reported for the DG at different ages together with no change in the number of stained astrocytes (Olabarria et al., 2010). Effects of normal aging on astrocytes in mice were also reported to be highly different between brain regions (i.e., GFAP-derived hypertrophy with age in hippocampal regions but atrophy in entorhinal cortex; Rodríguez et al., 2014). Our findings are consistent with this brain region-dependency, and emphasize the need to use various markers to interrogate different subpopulations of astrocytes. Globally, we report a higher number of cells expressing GFAP and ALDH1L1 in hippocampal regions and LH/thalamus, respectively, with an absence of detectable change in GFAP-derived morphology in 3xTg-AD females. Furthermore, we found the inhibition of SCD to result in CA1 astrocyte hypertrophy (i.e., increased radius), but to reduce the number of CA1 cells expressing GFAP. Fatty acids could be part of the mechanism by which SCDi reduces GFAP-positive cell count in CA1 in the studied AD model. Astrocytes were shown to respond to neuron-derived fatty acids by forming lipid droplets (Ioannou et al., 2019), and 3xTg-AD mice have been reported to have an increased number of lipid droplets in the subventricular zone (Hamilton et al., 2015), which could be associated with a higher metabolic demand on astrocytes. Since the inhibition of SCD was found to reduce lipids, specifically AD-associated triglycerides, in 3xTg-AD mice (Hamilton et al., 2015), it could contribute to a reduction in the need for metabolically active astrocytes to deal with lipids, and consequently to the lowered number of GFAP-expressing cells. Lipid-mediated dysregulation of astrocytic function could in parallel impact their regulatory role in synaptic plasticity (Murphy-Royal et al., 2020), which is relevant to the regulation of wake/sleep states and could also be rescued by inhibiting SCD.

A first limitation of the present study concerns the use of female mice only. This choice was guided by the more severe Aβ pathology reported in female 3xTg mice in comparison to males (Hirata-Fukae et al., 2008), and by the higher risk of developing AD in women (Lautenschlager et al., 1996). However, recent data indicates a higher efficiency of AD-targeting treatments in men in comparison to women (Kurkinen, 2023), and future investigations of sleep after SCD inhibition or related interventions should include males. Secondly, even if justified from previous research (Hamilton et al., 2015, 2022), the present study has solely evaluated the impact of a 28-day treatment administered at a relatively early age (approximately 4 months). It is possible that longer treatments as well as treatment targeting an age range with milder wake/sleep alterations could have resulted in some level of rescue of wake/sleep variables. In addition, a single concentration of SCDi was used in the present study (i.e., 80 μM) and, although within the range of doses shown to have positive effects on AD-related phenotypes in 3xTg mice (Hamilton et al., 2015, 2022), testing the efficacy of a range of doses in the context of wake/sleep evaluation would be an important next step. Lastly, ECoG activity was interrogated for the motor and visual cortex while astrocytes were examined in the hippocampus, LH, and thalamic nuclei. This methodological choice was made to simultaneously capture sleep regulatory regions (e.g., LH) and the hippocampus, because of the previously described modifications of hippocampal astrocytes in 3xTg-AD mice (Olabarria et al., 2010, 2011), but future work should interrogate astrocyte functioning at the site of ECoG recording.

## CONCLUSIONS

The present dataset highlights an extensive range of modifications in wake/sleep architecture and ECoG quality in a widely used mouse model of AD. This includes the original finding of higher multifractal complexity during wakefulness and SWS in (relatively young) 3xTg-AD mice, which has a high relevance to the understanding of mechanisms underlying the development of impaired cognitive functioning in neurodegenerative diseases (Averna et al., 2023). These changes in wake/sleep phenotypes were not rescued by the inhibition of SCD. Nonetheless, we have identified a vigilance state-specific ECoG signature of SCD inhibition, an observation pointing to a role of lipid metabolism in the regulation of synchronized cerebral cortex activity during wakefulness and sleep. SCD inhibition was previously found to reduce the activation of microglia in the 3xTg-AD model (Hamilton et al., 2022), and in combination with our current findings regarding the astrocytic marker GFAP, these discoveries suggest that glial populations are important targets of this intervention. Future research could evaluate how the different astrocytic functions, assembled recently in a framework as a contextual guide/integrator (Murphy-Royal et al., 2023), are contributing to lipid-dependent changes in wake/sleep quality using, for instance, manipulations of specific lipid metabolism-related enzymes in astrocytes.

## DECLARATIONS

### Authors’ contributions

A.H., K.F., J.B., and V.M. designed the experiments and analytic plan. A.H. and J.D.-G. performed the animal experiments. A.H. and T.L. analyzed the electrophysiological data. A.H. and M.J.d.C.C. conducted IHC experiments and analyses. B.D.-L. provided assistance with IHC. A.H., M.J.d.C.C., and T.L. performed statistical analyses and graphical representations. C.B. and J.-M.L. produced the code for scale-free activity analysis. A.H., M.J.d.C.C., and V.M. wrote and revised the manuscript. V.M. supervised all experiments and analyses. A.H., M.J.d.C.C., T.L., K.F., J.B., and V.M. provided funding. All authors approved the final version of the manuscript.

### Data availability

All data are included in this article and its supporting files. Raw data are available from the corresponding author upon reasonable request.

## Acknowledgements

The authors are thankful to Laura Hamilton, Lewis Rhys Depaauw-Holt and Ciaran Murphy-Royal for help and advice related to experiment and interpretation. The authors also want to thank Aurélie Cleret-Buhot from the cellular imaging platform of the Centre de recherche du Centre hospitalier de l’Université de Montréal (CRCHUM) for precious help with the development of the microscopy pipeline, and Marie-Claude Therrien from the animal facility (CRCHUM) for assistance with animal transfer.

## ABBREVIATIONS

3xTg: triple transgenic
aCSF: artificial cerebrospinal fluid
AD: Alzheimer’s disease
ALDH1L1: aldehyde dehydrogenase 1 family member L1 or 10-formyltetrahydrofolate dehydrogenase
ANOVA: analysis of variance
DAPI: 4′,6-diamidino-2-phenylindole
DG: dentate gyrus
DMSO: dimethylsulfoxide
ECoG: electrocorticographic
EMG: electromyographic
GFAP: glial fibrillary acidic protein
IHC: immunohistochemistry
LH: lateral hypothalamus
PBS: phosphate-buffered saline
PS: paradoxical sleep
SCD: stearoyl-CoA desaturase
SCDi: stearoyl-CoA desaturase inhibitor
SEM: standard error of the mean
SWA: slow wave activity
SWS: slow wave sleep
WT: wild-type.

## REFERENCES

Adler P, Mayne J, Walker K, Ning Z, Figeys D. 2019. Therapeutic targeting of casein kinase 1δ/ε in an Alzheimer’s disease mouse model. J. Proteome Res. 18, 3383–3393. 10.1021/acs.jproteome.9b00312.

Areal CC, Cao R, Sonenberg N, Mongrain V. 2020. Wakefulness/sleep architecture and electroencephalographic activity in mice lacking the translational repressor 4E-BP1 or 4E-BP2. Sleep 43, zsz210. 10.1093/sleep/zsz210.

Arsenault D, Julien C, Tremblay C, Calon F. 2011. DHA improves cognition and prevents dysfunction of entorhinal cortex neurons in 3xTg-AD mice. PLoS One 6, e17397. 10.1371/journal.pone.0017397.

Astarita G, Jung KM, Vasilevko V, Dipatrizio NV, Martin SK, Cribbs DH, Head E, Cotman CW, Piomelli D. 2011. Elevated stearoyl-CoA desaturase in brains of patients with Alzheimer’s disease. PLoS One 6, e24777. 10.1371/journal.pone.0024777.

Averna A, Coelli S, Ferrara R, Cerutti S, Priori A, Bianchi AM. 2023. Entropy and fractal analysis of brain-related neurophysiological signals in Alzheimer’s and Parkinson’s disease. J. Neural Eng. 20, 051001. 10.1088/1741-2552/acf8fa.

Ballester Roig MN, Leduc T, Dufort-Gervais J, Maghmoul Y, Tastet O, Mongrain V. 2023. Probing pathways by which rhynchophylline modifies sleep using spatial transcriptomics. Biol. Direct. 18, 21. 10.1186/s13062-023-00377-7.

Barki N, Bolognini D, Börjesson U, Jenkins L, Riddell J, Hughes DI, Ulven T, Hudson BD, Ulven ER, Dekker N, Tobin AB, Milligan G. 2022. Chemogenetics defines a short-chain fatty acid receptor gut-brain axis. Elife. 11, e73777. 10.7554/eLife.73777.

Belfiore R, Rodin A, Ferreira E, Velazquez R, Branca C, Caccamo A, Oddo S. 2019. Temporal and regional progression of Alzheimer’s disease-like pathology in 3xTg-AD mice. Aging Cell. 18, e12873. 10.1111/acel.12873.

Bell LN, Kilkus JM, Booth JN 3rd, Bromley LE, Imperial JG, Penev PD. 2013. Effects of sleep restriction on the human plasma metabolome. Physiol. Behav. 122, 25–31. 10.1016/j.physbeh.2013.08.007.

Benthem SD, Skelin I, Moseley SC, Stimmell AC, Dixon JR, Melilli AS, Molina L, McNaughton BL, Wilber AA. 2020. Impaired hippocampal-cortical interactions during sleep in a mouse model of Alzheimer’s disease. Curr. Biol. 30, 2588–2601.e5. 10.1016/j.cub.2020.04.087.

Broussard JL, Chapotot F, Abraham V, Day A, Delebecque F, Whitmore HR, Tasali E. 2015. Sleep restriction increases free fatty acids in healthy men. Diabetologia 58, 791–798. 10.1007/s00125-015-3500-4.

Cai P, Huang SN, Lin ZH, Wang Z, Liu RF, Xiao WH, Li ZS, Zhu ZH, Yao J, Yan XB, Wang FD, Zeng SX, Chen GQ, Yang LY, Sun YK, Yu C, Chen L, Wang WX. 2022. Regulation of wakefulness by astrocytes in the lateral hypothalamus. Neuropharmacology 221, 109275. 10.1016/j.neuropharm.2022.109275.

Chan RB, Oliveira TG, Cortes EP, Honig LS, Duff KE, Small SA, Wenk MR, Shui G, Di Paolo G. 2012. Comparative lipidomic analysis of mouse and human brain with Alzheimer disease. J. Biol. Chem. 287, 2678–2688. 10.1074/jbc.M111.274142.

Chatterjee P, Lim WL, Shui G, Gupta VB, James I, Fagan AM, Xiong C, Sohrabi HR, Taddei K, Brown BM, Benzinger T, Masters C, Snowden SG, Wenk MR, Bateman RJ, Morris JC, Martins RN. 2016. Plasma phospholipid and sphingolipid alterations in Presenilin1 mutation carriers: a pilot study. J. Alzheimers Dis. 50, 887–894. 10.3233/JAD-150948.

Chen ZP, Wang S, Zhao X, Fang W, Wang Z, Ye H, Wang MJ, Ke L, Huang T, Lv P, Jiang X, Zhang Q, Li L, Xie ST, Zhu JN, Hang C, Chen D, Liu X, Yan C. 2023. Lipid-accumulated reactive astrocytes promote disease progression in epilepsy. Nat. Neurosci. 26, 542–554. 10.1038/s41593-023-01288-6.

Ciuciu P, Abry P, Rabrait C, Wendt H. 2008. Log-wavelet Leaders cumulant based multifractal analysis of EVI fMRI time series: Evidence of scaling in ongoing and evoked brain activity. IEEE J. Sel. Top. Signal Process. 2, 929–943. 10.1109/JSTSP.2008.2006663.

Clasadonte J, Scemes E, Wang Z, Boison D, Haydon PG. 2017. Connexin 43-mediated astroglial metabolic networks contribute to the regulation of the sleep-wake cycle. Neuron 95, 1365–1380.e5. 10.1016/j.neuron.2017.08.022.

Dufort-Gervais J, Mongrain V, Brouillette J. 2019. Bidirectional relationships between sleep and amyloid-beta in the hippocampus. Neurobiol. Learn. Mem. 160, 108–117. 10.1016/j.nlm.2018.06.009.

El Helou J, Bélanger-Nelson E, Freyburger M, Dorsaz S, Curie T, La Spada F, Gaudreault PO, Beaumont É, Pouliot P, Lesage F, Frank MG, Franken P, Mongrain V. 2013. Neuroligin-1 links neuronal activity to sleep-wake regulation. Proc. Natl. Acad. Sci. USA 110, 9974–9979. 10.1073/pnas.1221381110.

Foley J, Blutstein T, Lee S, Erneux C, Halassa MM, Haydon P. 2017. Astrocytic IP3/Ca2+ signaling modulates theta rhythm and REM sleep. Front. Neural Circuits. 11, 3. 10.3389/fncir.2017.00003.

Fraser T, Tayler H, Love S. 2010. Fatty acid composition of frontal, temporal and parietal neocortex in the normal human brain and in Alzheimer’s disease. Neurochem. Res. 35, 503–513. 10.1007/s11064-009-0087-5.

González-Molina LA, Villar-Vesga J, Henao-Restrepo J, Villegas A, Lopera F, Cardona-Gómez GP, Posada-Duque R. 2021. Extracellular vesicles from 3xTg-AD mouse and Alzheimer’s disease patient astrocytes impair neuroglial and vascular components. Front. Aging Neurosci. 13, 593927. 10.3389/fnagi.2021.593927.

Halassa MM, Florian C, Fellin T, Munoz JR, Lee SY, Abel T, Haydon PG, Frank MG. 2009. Astrocytic modulation of sleep homeostasis and cognitive consequences of sleep loss. Neuron 61, 213–219. 10.1016/j.neuron.2008.11.024.

Hamilton LK, Dufresne M, Joppé SE, Petryszyn S, Aumont A, Calon F, Barnabé-Heider F, Furtos A, Parent M, Chaurand P, Fernandes KJ. 2015. Aberrant lipid metabolism in the forebrain niche suppresses adult neural stem cell proliferation in an animal model of Alzheimer’s disease. Cell. Stem. Cell. 17, 397–411. 10.1016/j.stem.2015.08.001.

Hamilton LK, Moquin-Beaudry G, Mangahas CL, Pratesi F, Aubin M, Aumont A, Joppé SE, Légiot A, Vachon A, Plourde M, Mounier C, Tétreault M, Fernandes KJL. 2022. Stearoyl-CoA Desaturase inhibition reverses immune, synaptic and cognitive impairments in an Alzheimer’s disease mouse model. Nat. Commun. 13, 2061. 10.1038/s41467-022-29506-y. Erratum in: 10.1038/s41467-023-38295-x.

Hector A, Provost C, Delignat-Lavaud B, Bouamira K, Menaouar CA, Mongrain V, Brouillette J. 2023 Hippocampal injections of soluble amyloid-beta oligomers alter electroencephalographic activity during wake and slow wave sleep in rats. Alzheimers Res. Ther. 15, 174. 10.1186/s13195-023-01316-4.

Heneka MT, Carson MJ, El Khoury J, Landreth GE, Brosseron F, Feinstein DL, Jacobs AH, Wyss-Coray T, Vitorica J, Ransohoff RM, Herrup K, Frautschy SA, Finsen B, Brown GC, Verkhratsky A, Yamanaka K, Koistinaho J, Latz E, Halle A, Petzold GC, Town T, Morgan D, Shinohara ML, Perry VH, Holmes C, Bazan NG, Brooks DJ, Hunot S, Joseph B, Deigendesch N, Garaschuk O, Boddeke E, Dinarello CA, Breitner JC, Cole GM, Golenbock DT, Kummer MP. 2015. Neuroinflammation in Alzheimer’s disease. Lancet Neurol. 14, 388–405. 10.1016/S1474-4422(15)70016-5.

Hirata-Fukae C, Li HF, Hoe HS, Gray AJ, Minami SS, Hamada K, Niikura T, Hua F, Tsukagoshi-Nagai H, Horikoshi-Sakuraba Y, Mughal M, Rebeck GW, LaFerla FM, Mattson MP, Iwata N, Saido TC, Klein WL, Duff KE, Aisen PS, Matsuoka Y. 2008. Females exhibit more extensive amyloid, but not tau, pathology in an Alzheimer transgenic model. Brain Res. 1216, 92–103. 10.1016/j.brainres.2008.03.079.

Ioannou MS, Jackson J, Sheu SH, Chang CL, Weigel AV, Liu H, Pasolli HA, Xu CS, Pang S, Matthies D, Hess HF, Lippincott-Schwartz J, Liu Z. 2019. Neuron-astrocyte metabolic coupling protects against activity-induced fatty acid toxicity. Cell. 177, 1522–1535.e14. doi: 10.1016/j.cell.2019.04.001.

Kent BA, Feldman HH, Nygaard HB. 2021. Sleep and its regulation: An emerging pathogenic and treatment frontier in Alzheimer’s disease. Prog. Neurobiol. 197, 101902. 10.1016/j.pneurobio.2020.101902.

Kent BA, Michalik M, Marchant EG, Yau KW, Feldman HH, Mistlberger RE, Nygaard HB. 2019. Delayed daily activity and reduced NREM slow-wave power in the APPswe/PS1dE9 mouse model of Alzheimer’s disease. Neurobiol. Aging. 78, 74–86. 10.1016/j.neurobiolaging.2019.01.010.

Kent BA, Strittmatter SM, Nygaard HB. 2018. Sleep and EEG power spectral analysis in three transgenic mouse models of Alzheimer’s disease: APP/PS1, 3xTgAD, and Tg2576. J. Alzheimers Dis. 64, 1325–1336. 10.3233/JAD-180260.

Klein M, Lohr C, Droste D. 2020. Age-dependent heterogeneity of murine olfactory bulb astrocytes.

Front. Aging Neurosci. 12, 172. 10.3389/fnagi.2020.00172.

Kosel F, Pelley JMS, Franklin TB. 2020. Behavioural and psychological symptoms of dementia in mouse models of Alzheimer’s disease-related pathology. Neurosci. Biobehav. Rev. 112, 634–647. 10.1016/j.neubiorev.2020.02.012.

Kurkinen M. 2023. Lecanemab (Leqembi) is not the right drug for patients with Alzheimer’s disease. Adv. Clin. Exp. Med. 32, 943–947. 10.17219/acem/171379.

Lau ZJ, Pham T, Chen SHA, Makowski D. 2022. Brain entropy, fractal dimensions and predictability: A review of complexity measures for EEG in healthy and neuropsychiatric populations. Eur. J. Neurosci. 56, 5047–5069. 10.1111/ejn.15800.

Lautenschlager NT, Cupples LA, Rao VS, Auerbach SA, Becker R, Burke J, Chui H, Duara R, Foley EJ, Glatt SL, Green RC, Jones R, Karlinsky H, Kukull WA, Kurz A, Larson EB, Martelli K, Sadovnick AD, Volicer L, Waring SC, Growdon JH, Farrer LA. 1996. Risk of dementia among relatives of Alzheimer’s disease patients in the MIRAGE study: what is in store for the oldest old? Neurology. 46, 641–650. 10.1212/WNL.46.3.641.

Lee JA, Hall B, Allsop J, Alqarni R, Allen SP. 2021. Lipid metabolism in astrocytic structure and function. Semin. Cell Dev. Biol. 112, 123–136. 10.1016/j.semcdb.2020.07.017.

Lee YF, Russ AN, Zhao Q, Perle SJ, Maci M, Miller MR, Hou SS, Algamal M, Zhao Z, Li H, Gelwan N, Liu Z, Gomperts SN, Araque A, Galea E, Bacskai BJ, Kastanenka KV. 2023. Optogenetic targeting of astrocytes restores slow brain rhythm function and slows Alzheimer’s disease pathology. Sci. Rep. 13, 13075.

Lina JM, O’Callaghan EK, Mongrain V. 2019. Scale-Free Dynamics of the mouse wakefulness and sleep electroencephalogram quantified using Wavelet-Leaders. Clocks Sleep. 1, 50–64. 10.3390/clockssleep1010006.

Ma Q, Ning X, Wang J, Li J. 2005. Sleep-stage characterization by nonlinear EEG analysis using Wavelet-based multifractal formalism. Conf. Proc. IEEE Eng. Med. Biol. Soc. 2005, 4526–4529. 10.1109/IEMBS.2005.1615475.

Mi Y, Qi G, Vitali F, Shang Y, Raikes AC, Wang T, Jin Y, Brinton RD, Gu H, Yin F. 2023. Loss of fatty acid degradation by astrocytic mitochondria triggers neuroinflammation and neurodegeneration. Nat. Metab. 5, 445–465. 10.1038/s42255-023-00756-4.

Murphy-Royal C, Ching S, Papouin T. 2023. A conceptual framework for astrocyte function. Nat. Neurosci. 26, 1848–1856. 10.1038/s41593-023-01448-8.

Murphy-Royal C, Johnston AD, Boyce AKJ, Diaz-Castro B, Institoris A, Peringod G, Zhang O, Stout RF, Spray DC, Thompson RJ, Khakh BS, Bains JS, Gordon GR. 2020. Stress gates an astrocytic energy reservoir to impair synaptic plasticity. Nat. Commun. 11, 2014. 10.1038/s41467-020-15778-9.

Oddo S, Caccamo A, Shepherd JD, Murphy MP, Golde TE, Kayed R, Metherate R, Mattson MP, Akbari Y, LaFerla FM. 2003. Triple-transgenic model of Alzheimer’s disease with plaques and tangles: intracellular Abeta and synaptic dysfunction. Neuron. 39, 409–421. 10.1016/S0896-6273(03)00434-3.

Olabarria M, Noristani HN, Verkhratsky A, Rodríguez JJ. 2010. Concomitant astroglial atrophy and astrogliosis in a triple transgenic animal model of Alzheimer’s disease. Glia 58, 831–838. 10.1002/glia.20967.

Olabarria M, Noristani HN, Verkhratsky A, Rodríguez JJ. 2011. Age-dependent decrease in glutamine synthetase expression in the hippocampal astroglia of the triple transgenic Alzheimer’s disease mouse model: mechanism for deficient glutamatergic transmission? Mol. Neurodegener. 6, 55. 10.1186/1750-1326-6-55.

Rodríguez JJ, Yeh CY, Terzieva S, Olabarria M, Kulijewicz-Nawrot M, Verkhratsky A. 2014. Complex and region-specific changes in astroglial markers in the aging brain. Neurobiol. Aging 35, 15–23. 10.1016/j.neurobiolaging.2013.07.002.

Saber M, Murphy SM, Cho Y, Lifshitz J, Rowe RK. 2021. Experimental diffuse brain injury and a model of Alzheimer’s disease exhibit disease-specific changes in sleep and incongruous peripheral inflammation. J. Neurosci. Res. 99, 1136–1160. 10.1002/jnr.24771.

Shi L, Chen SJ, Ma MY, Bao YP, Han Y, Wang YM, Shi J, Vitiello MV, Lu L. 2018. Sleep disturbances increase the risk of dementia: A systematic review and meta-analysis. Sleep Med. Rev. 40, 4–16. 10.1016/j.smrv.2017.06.010.

Skuladottir GV, Nilsson EK, Mwinyi J, Schiöth HB. 2016. One-night sleep deprivation induces changes in the DNA methylation and serum activity indices of stearoyl-CoA desaturase in young healthy men. Lipids Health Dis. 15, 137. 10.1186/s12944-016-0309-1.

Smolič T, Tavčar P, Horvat A, Černe U, Halužan Vasle A, Tratnjek L, Kreft ME, Scholz N, Matis M, Petan T, Zorec R, Vardjan N. 2021. Astrocytes in stress accumulate lipid droplets. Glia 69, 1540–1562. 10.1002/glia.23978.

Sterniczuk R, Dyck RH, Laferla FM, Antle MC. 2010. Characterization of the 3xTg-AD mouse model of Alzheimer’s disease: part 1. Circadian changes. Brain Res. 1348, 139–148. 10.1016/j.brainres.2010.05.013.

Tavares, G., Martins, M., Correia, J.S., Sardinha, V.M., Guerra-Gomes, S., das Neves, S.P., Marques, F., Sousa, N., Oliveira, J.F. 2017. Employing an open-source tool to assess astrocyte tridimensional structure. Brain Struct. Funct. 222, 1989–1999. 10.1007/s00429-016-1316-8.

Vassalli A, Franken P. 2017. Hypocretin (orexin) is critical in sustaining theta/gamma-rich waking behaviors that drive sleep need. Proc. Natl. Acad. Sci. USA. 114, E5464–E5473. 10.1073/pnas.1700983114.

Weiss B, Clemens Z, Bódizs R, Vágó Z, Halász P. 2009. Spatio-temporal analysis of monofractal and multifractal properties of the human sleep EEG. J. Neurosci. Methods 185, 116–124. 10.1016/j.jneumeth.2009.07.027.

Whiley L, Sen A, Heaton J, Proitsi P, García-Gómez D, Leung R, Smith N, Thambisetty M, Kloszewska I, Mecocci P, Soininen H, Tsolaki M, Vellas B, Lovestone S, Legido-Quigley C; AddNeuroMed Consortium. 2014. Evidence of altered phosphatidylcholine metabolism in Alzheimer’s disease. Neurobiol. Aging. 35, 271–278. 10.1016/j.neurobiolaging.2013.08.001.

Wang C, Holtzman DM. 2020. Bidirectional relationship between sleep and Alzheimer’s disease: role of amyloid, tau, and other factors. Neuropsychopharmacology 45, 104–120. 10.1038/s41386-019-0478-5.

Wendt H, Roux SG, Abry P, Jaffard S. 2008. Wavelet-leaders and bootstrap for multifractal analysis of images. Signal Process. 89, 1100–1114. 10.1016/j.sigpro.2008.12.015.

Wu R, Tripathy S, Menon V, Yu L, Buchman AS, Bennett DA, De Jager PL, Lim ASP. 2023. Fragmentation of rest periods, astrocyte activation, and cognitive decline in older adults with and without Alzheimer’s disease. Alzheimers Dement. 19, 1888–1900. 10.1002/alz.12817.

Yeh CY, Vadhwana B, Verkhratsky A, Rodríguez JJ. 2011. Early astrocytic atrophy in the entorhinal cortex of a triple transgenic animal model of Alzheimer’s disease. ASN Neuro. 3, 271–279. 10.1042/AN20110025.

Zhang Y, Ren R, Yang L, Zhang H, Shi Y, Okhravi HR, Vitiello MV, Sanford LD, Tang X. 2022. Sleep in Alzheimer’s disease: a systematic review and meta-analysis of polysomnographic findings. Transl. Psychiatry. 12, 136. 10.1038/s41398-022-01897-y.

